# MicroRNAs reinforce repression of PRC2 transcriptional targets independently and through a feed-forward regulatory network

**DOI:** 10.1101/298919

**Authors:** Haridha Shivram, Steven V. Le, Vishwanath R. Iyer

**Affiliations:** Center for Systems and Synthetic Biology, Institute for Cellular and Molecular Biology, Department of Molecular Biosciences, University of Texas at Austin, Austin, Texas 78712, United States of America

**Keywords:** EZH2, Polycomb repressive complex (PRC2), microRNA, AGO2, *Sleeping Beauty* transposon system, Vaccinia virus, VP55, RNA-binding, feed-forward, regulatory network, iCLIP, epigenetics

## Abstract

Gene expression can be regulated at multiple levels, but it is not known if and how there is broad coordination between regulation at the transcriptional and post-transcriptional levels. Transcription factors and chromatin regulate gene expression transcriptionally, while microRNAs (miRNAs) are small regulatory RNAs that function post-transcriptionally. Systematically identifying the post-transcriptional targets of miRNAs and the mechanism of transcriptional regulation of the same targets can shed light on regulatory networks connecting transcriptional and post-transcriptional control. We used iCLIP (individual crosslinking and immunoprecipitation) for the RISC (RNA-induced silencing complex) component AGO2 and global miRNA depletion to identify genes directly targeted by miRNAs. We found that PRC2 (Polycomb repressive complex 2) and its associated histone mark, H3K27me3, is enriched at hundreds of miRNA-repressed genes. We show that these genes are directly repressed by PRC2 and constitute a significant proportion of direct PRC2 targets. For just over half of the genes co-repressed by PRC2 and miRNAs, PRC2 promotes their miRNA-mediated repression by increasing expression of the miRNAs that are likely to target them. miRNAs also repress the remainder of the PRC2 target genes, but independently of PRC2. Thus, miRNAs post-transcriptionally reinforce silencing of PRC2-repressed genes that are inefficiently repressed at the level of chromatin, by either forming a feed-forward regulatory network with PRC2 or repressing them independently of PRC2.

## Introduction

Transcription factors (TFs) and miRNAs together form the largest components of gene regulatory networks and can regulate gene expression through both distinct and coordinated regulatory mechanisms. One of the ways in which TFs can coordinate their regulatory impact with miRNAs is by forming a feed-forward regulatory network. In such a network, a TF regulates a miRNA and both co-regulate a common target. In a coherent feed-forward regulatory network, the outcomes of both direct and indirect regulation by a TF are consistent. Many such TF-miRNA feed-forward regulatory networks have been shown to functionally impact several processes in development and disease (O’Donnell et al. 2005; Hobert 2008; Tsang et al. 2007; Lin et al. 2015; Gerloff et al. 2014; Polioudakis et al. 2013).

One potential function of coherent TF-miRNA feed-forward regulatory networks is to reinforce transcriptional regulation at the post-transcriptional level. In particular, this can help suppress residual transcripts produced from leaky transcription of transcriptionally silenced genes. This is most crucial during switches in transcriptional states in response to stress, developmental transitions, cell cycle stages or other external stimuli (Farh et al. 2005; WU et al. 2015; del Rosario et al. 2016).

PRC2 is an epigenetic regulator complex that transcriptionally silences genes through modification of histone H3 with tri-methylation at lysine 27 (H3K27me3). PRC2 plays a critical role in maintaining the silenced state of genes involved in development and several cancers (Aranda et al. 2015; Di Croce and Helin 2013). In this study, we provide data that points to a broad role of miRNAs where they independently strengthen and also reinforce the silencing of PRC2-repressed genes post-transcriptionally.

## Results

### Transcriptome-wide identification of miRNA targets

miRNAs primarily target mRNAs through interactions between their 5’ seed region and the 3’ untranslated region (3’ UTR) of the mRNA, mediated by the RNA-induced silencing complex (RISC). To identify the transcriptome-wide targets of miRNAs, we first performed RNA individual crosslinking and immunoprecipitation (iCLIP) for the RISC component AGO2 in glioblastoma multiforme (GBM) cells (König et al. 2010). Several observations indicate that the AGO2-RNA interactions we identified using iCLIP were direct and specific. First, AGO2-RNA complexes were crosslinking- and RNase-treatment-specific (Fig. 1A). Second, reads from iCLIP were enriched at 3’ UTRs as expected from the binding of RISC (Fig. 1B). Third, reads mapping to genes showed high correlation and were reproducible across independent biological replicate experiments (Fig. 1C). We identified 45362 peaks mapping to 5896 protein-coding genes that were common across two independent biological replicate experiments.

**Figure 1.**
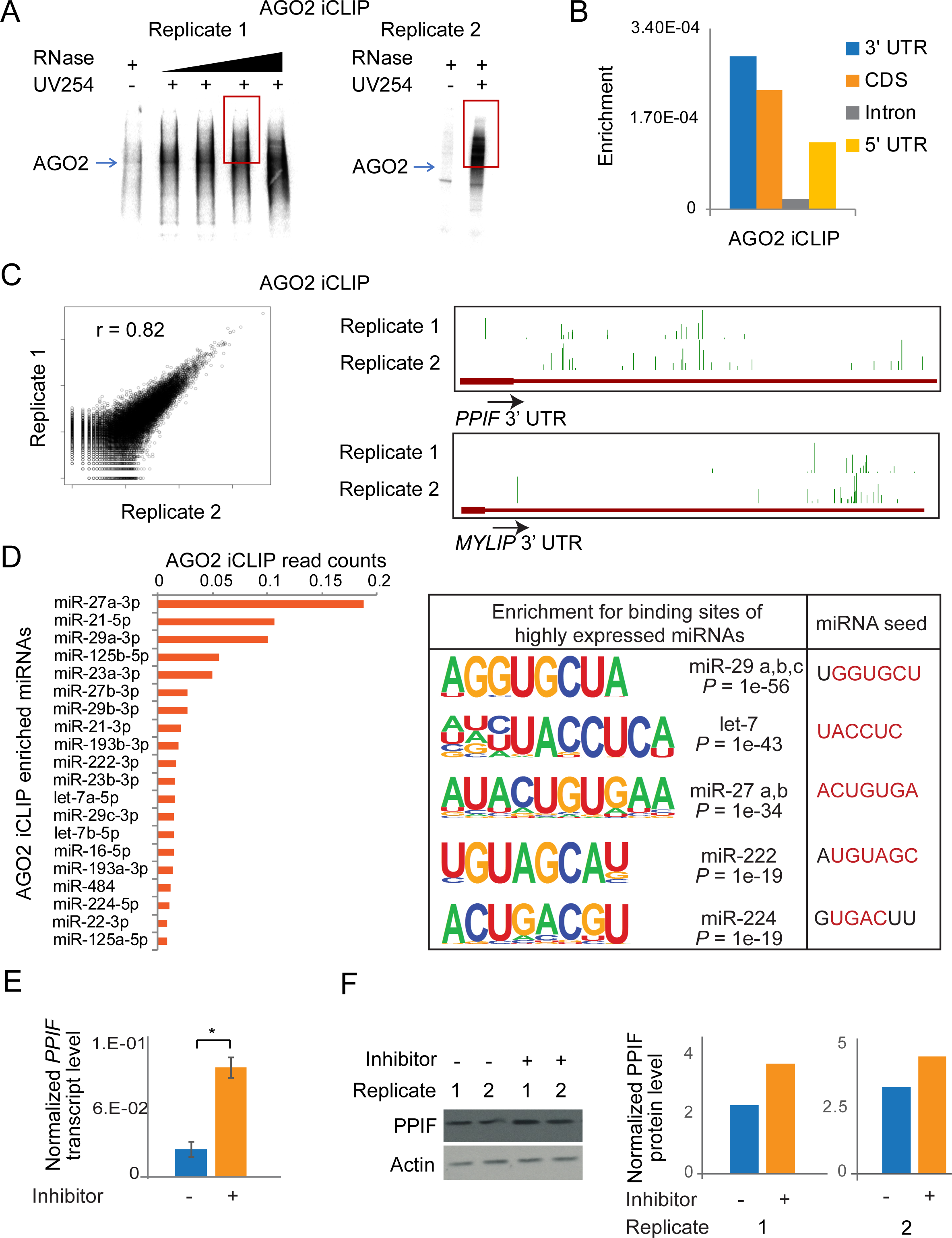
Identification of miRNA targets by AGO2 iCLIP. (A) Autoradiograph showing AGO2-crosslinked ribonucleoprotein complexes from 2 replicates. The red box represents the region that was processed to make cDNA libraries. (B) Genomic enrichment of AGO2-crosslinked reads mapping to mRNAs. The plot shows total reads mapped to a genomic feature, normalized to the total length of the feature in the genome. (C) Correlation of the total reads mapping to genes between the two independent replicates of AGO2 iCLIP (left panel). Genome browser images showing AGO2 iCLIP peaks at 3’ UTRs for two protein-coding genes (right panel). (D) AGO2-interacting miRNAs plotted in descending order of read coverage (left panel). Sequences enriched proximal to AGO2 iCLIP peaks on mRNAs that significantly match the binding sequence for AGO2-interacting miRNAs. Corresponding miRNA seed sequences are also shown (right panel). (E) *PPIF* transcript levels in U87MG cells treated with negative control inhibitor or *miR-23a-3p* inhibitor. Error bars represent standard error across 4 replicates (2 biological and 2 technical). * indicates *P* < 0.05 (paired t-test). (F) PPIF protein levels in U87MG cells treated with negative control inhibitor or *miR-23a-3p* inhibitor. The immunoblot showing two independent replicates (left panel) was quantified using ImageJ (right panel).

In addition to mRNA targets, the AGO2 iCLIP experiment also recovered miRNAs that AGO2 interacts with, allowing for more detailed analysis of functional miRNA-mRNA interactions. Although enrichment of miRNAs in AGO2 iCLIP showed high correlation to their expression levels, miRNAs with the highest expression were not necessarily the most enriched (Supplemental Fig. S1A). We found that the region spanning 200 bases around AGO2 iCLIP peaks was significantly enriched for the seed sequences of several of the top AGO2-interacting miRNA families (Fig. 1D and Supplemental Fig. S1A). In addition, the predicted mRNA target sites of the top 10 highly enriched miRNAs were significantly more likely to occur proximal to AGO2 iCLIP peaks than background mRNAs (Supplemental Fig. S1B). This suggests that the miRNAs associated with AGO2 bind to a significant proportion of the AGO2-enriched mRNAs, further attesting to the ability of the iCLIP experiment to broadly identify the targets of active miRNAs.

To further support the miRNA-mRNA interactions we identified, we validated the regulation of *PPIF* by miR-23a-3p. miR-23a-3p was among the most enriched miRNAs in our AGO2 iCLIP dataset. We selected *PPIF* as a target of *miR-23a-3p* as it had a predicted binding site close to an AGO2 iCLIP peak (Supplemental Fig. S2A). Several lines of evidence show that miR-23a-3p directly represses *PPIF* expression. First, treatment of cells with an inhibitor of miR-23a-3p led to an induction of both *PPIF* transcript and protein levels (Fig. 1E,F). Second, we checked if the repression of *PPIF* was through a direct interaction between the 3’ UTR of *PPIF* and the miR-23a-3p seed region. We performed luciferase reporter assays where we cloned the 3’ UTR of *PPIF* and compared luciferase expression to constructs that were mutated at the sites of interaction. We found that the effect of inhibitor treatment on regulation of the *PPIF* 3’ UTR was abolished upon deletion of the miR-23a-3p binding sites (Supplemental Fig. S2B). Third, both deletion of the miRNA binding sites and disruption of miRNA-mRNA interactions through base substitutions led to a similar rescue in luciferase activity (Supplemental Fig. S2C).

### miRNA-repressed genes are enriched for PRC2 binding and H3K27me3

Although AGO2 iCLIP identified all the interaction targets of RISC, the AGO2-RISC complex can regulate gene expression in both a miRNA-dependent and independent manner (Leung et al. 2011). To specifically identify miRNA-dependent genes, we adopted a strategy to globally deplete miRNAs, followed by expression profiling. Ectopic overexpression of the Vaccinia Virus protein *VP55* has been previously shown to polyadenylate and degrade miRNAs, causing a global loss of miRNAs. Transcriptome changes in cells overexpressing *VP55* closely match that of cells in response to loss of DICER, indicating that *VP55* overexpression does not cause major transcriptional perturbations independent of its effect on miRNAs (Backes et al. 2012; Aguado et al. 2015). We cloned *VP55* into a *Sleeping Beauty* transposon system under a doxycycline-inducible promoter and stably integrated this construct into T98G cells to generate a GBM cell line capable of doxycycline-responsive loss of miRNAs (Kowarz et al. 2015). miR-21, the most abundant miRNA in these cells, was almost completely depleted within 24 hours of doxycycline treatment (Fig. 2A). PCR-amplified cDNA libraries prior to size-selection for miRNA-seq showed a significant depletion of miRNAs (Supplemental Fig. S3A). miRNA-seq showed a global depletion of approximately 90% of all cellular miRNAs (Fig. 2B). To identify the genes regulated by miRNAs genome-wide, we performed mRNA-seq from doxycycline treated (+ Dox) and untreated cells (-Dox). 3844 protein-coding transcripts were up-regulated and thus repressed by miRNAs, and 3668 transcripts were down-regulated upon loss of miRNAs (Supplemental Fig. S3B). Some of the most strongly miRNA-repressed genes included several genes from the histone gene family. This is consistent with *VP55*-responsive genes identified previously in HEK293T cells (Aguado et al. 2015).

**Figure 2.**
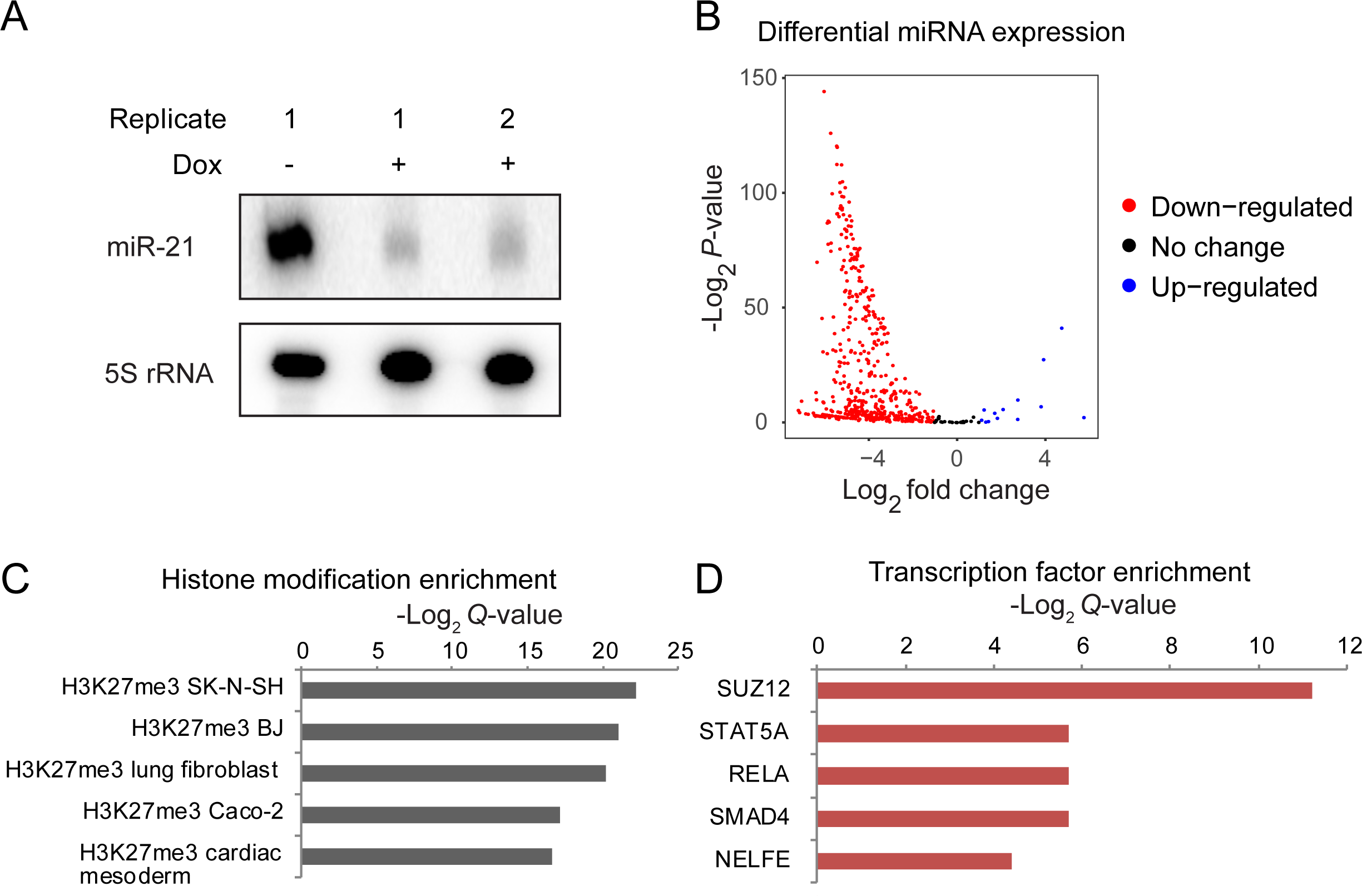
miRNA-repressed genes are enriched for PRC2 binding and H3K27me3. (A) miR-21 expression levels in response to *VP55* induction. Dox = Doxycycline. (B) Volcano plot of differentially expressed miRNAs in response to *VP55* induction. X-axis represents log_2_ fold difference between expression of miRNAs in *VP55*-induced and non-induced cells. Y-axis represents −log_2_ *P*-value of the expression difference calculated using DESeq2. (C,D) Enrichment for H3K27me3 (C) and SUZ12 (D) at genes derepressed in miRNA depleted cells (miRNA-repressed genes). Enrichments were calculated using Enrichr (Chen et al. 2013).

37% of the miRNA-repressed transcripts interacted directly with AGO2 based on iCLIP, suggesting that these were direct functional targets of miRNAs. Thus, miRNA-repressed genes include direct as well as indirect targets which could be regulated at the level of transcription (Gosline et al. 2016). To check if the miRNA-repressed genes were also regulated by specific transcription factors, we checked for the enrichment of TF binding sites proximal to their promoters. We used Enrichr to detect TF regulatory signatures in the ranked list of miRNA-repressed genes (Chen et al. 2013). We found a significant enrichment for H3K27me3 (Fig. 2C) and a corresponding enrichment for SUZ12, a member of the PRC2 complex which adds H3K27me3 (Fig. 2D), at miRNA-repressed genes. This suggested the possibility that many miRNA-repressed genes could additionally be regulated at the level of chromatin by PRC2 and H3K27 methylation. As a control, we also performed Enrichr analysis using a set of genes expressed in the same range as the miRNA-repressed genes, but not repressed by miRNAs. The control set showed no enrichment for any specific TF but did show an enrichment for H3K9me3 and H3K27me3, as would be expected from low-expressed genes (Supplemental Fig. S4A). To determine if this could be a general mode of dual regulatory control in other cell types, we analyzed data from mouse embryonic stem cells (mESCs) with a knockout of the miRNA-processing factor DICER, which also results in loss of miRNAs (Zheng et al. 2014). Similar to GBM cells, we found that genes repressed by miRNAs in mESCs also showed enrichment for H3K27me3 and PRC2 (Supplemental Fig. S4B,C).

### miRNAs repress hundreds of genes directly repressed by PRC2

Based on the above results, we hypothesized that many genes that are post-transcriptionally repressed by miRNAs are also transcriptionally repressed by PRC2. To test this hypothesis, we generated a CRISPR-Cas9 knockout of *EZH2* in GBM cells. H3K27me3 levels, as well as levels of SUZ12, were reduced in *EZH2* -/- cells, signifying loss of the PRC2 complex (Supplemental Fig. S5A) (Pasini et al. 2004). 1519 protein-coding genes were down-regulated and 1834 protein-coding genes were up-regulated (derepressed) upon loss of EZH2 (Fig. 3A). A significant number of the *EZH2*-repressed genes (444 genes, *P* = 3.5e-08 by hypergeometric test) were also repressed by miRNAs. Consistent with our Enrichr analysis, this data suggests that the genes repressed by EZH2 are also repressed by miRNAs. Alternatively, low-expressed genes could be regulated by miRNAs irrespective of them being repressed by PRC2. To test this alternative possibility, we utilized a bootstrapping approach (Methods) where we tested the overlap of the 3844 miRNA-repressed genes with random sets of 1834 genes expressed at similar levels to those of the EZH2-repressed genes. This bootstrap analysis showed that the number of genes co-repressed by EZH2 and miRNAs was indeed highly significant (*P* < 1e-4). Thus, the overlap we observed between EZH2 and miRNA-repressed genes is not driven by the fact that both sets of genes are low-expressed genes.

To determine if the genes co-repressed by *EZH2* and miRNAs were directly regulated by *EZH2*, we performed ChIP-seq for EZH2, as well as for H3K27me3. We detected EZH2 binding proximal to the TSS of 6132 genes where the level of H3K27me3 positively correlated with that of EZH2 binding (Supplemental Fig. S5B,C). Genes derepressed upon loss of *EZH2* (*EZH2*-repressed genes) showed EZH2 binding and H3K27me3 (Fig. 3B and Supplemental Fig. S6A). Consistent with previous studies, *EZH2*-repressed genes also showed occupancy by H3K4me3 (Fig. 3B and Supplemental Fig. S6A), with the enrichment for EZH2 and H3K27me3 showing a reciprocal relationship to H3K4me3 at *EZH2*-repressed genes (Fig. 3C) (Jadhav et al. 2016; Abou El Hassan et al. 2015). 959 of the 1834 *EZH2*-repressed genes had an EZH2 chromatin peak close to their transcription start site (TSS) and were thus likely directly repressed by *EZH2*.

**Figure 3.**
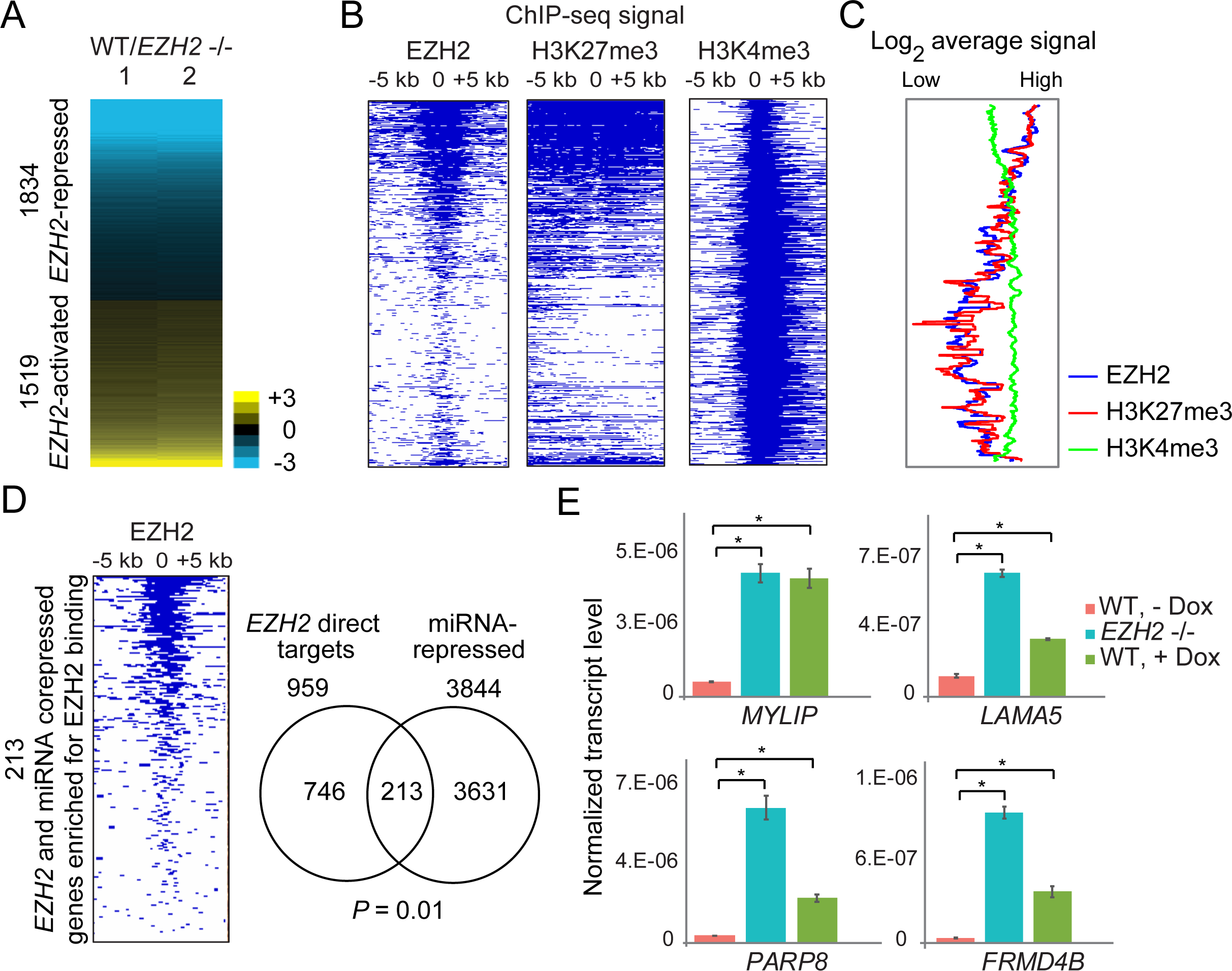
miRNAs repress hundreds of genes directly repressed by PRC2. (A) Heat map showing differential expression of protein-coding genes in response to *EZH2* knockout. Genes are ranked in increasing order of log_2_ fold change (≤ −0.5 and ≥ 0.5 log_2_ fold change) in WT/*EZH2* -/-. (B) Heat map showing enrichment scores (-log_10_ *Q*-value calculated using MACS2) for EZH2, H3K27me3 and H3K4me3 for genes plotted in and ordered as in A. (C) Average EZH2, H3K27me3 and H3K4me3 ChIP-seq signal for genes plotted in A. (D). Left: Heat map plotted as in B for genes co-repressed by *EZH2* and miRNAs. Right: Overlap between genes directly repressed by EZH2 and gene repressed by miRNAs. Significance of overlap was calculated using the hypergeometric test. (E) qRT-PCR validation of genes repressed by both *EZH2* and miRNAs, plotted as in Fig. 1E. * indicates *P* < 0.05 (paired t-test).

We then checked if the 444 genes co-repressed by *EZH2* and miRNAs were directly regulated by *EZH2*. Of these 444 genes, 213 genes showed EZH2 binding proximal to their TSS. These 213 genes that were jointly and directly repressed by *EZH2* and by miRNAs constituted 22% of all direct *EZH2* targets (*P* = 0.01 by hypergeometric test, Fig. 3D). This overlap was also significant (*P* = 0.0002) based on a bootstrapping approach to control for the low expression of the genes (Methods). We verified derepression upon loss of miRNAs or upon loss of *EZH2* of multiple genes using qRT-PCR (Fig. 3E). 34.7% of the genes co-repressed by *EZH2* and miRNAs also showed direct interaction with AGO2 by iCLIP (Supplemental Fig. S6B,C). miRNAs thus post-transcriptionally reinforce transcriptional repression by PRC2 of at least one-fifth of all PRC2 targets.

### miRNAs repress PRC2 target genes through a feed-forward regulatory network with PRC2

PRC2 and miRNAs were recently shown to independently co-repress endocytosis genes in mESCs (Mote et al. 2017). To test if miRNAs and PRC2 repressed a common set of genes independently in GBM cells, we depleted miRNAs globally by overexpressing *VP55* in *EZH2* -/- T98G cells and performed miRNA-seq and mRNA-seq. Most miRNAs were depleted within 24 hours of doxycycline treatment (Fig. 4A and Supplemental Fig. S7A). Both WT and *EZH2* -/- cells showed similar responses to doxycycline treatment in terms of the number of miRNAs depleted and the extent of depletion, indicating that *VP55* overexpression was equally effective in WT and *EZH2* -/- cells (Supplemental Fig. S7B,C). If miRNAs and *EZH2* repressed a common set of genes independently of one another, we would expect that these co-repressed genes would be repressed by miRNAs even in the absence of *EZH2*, and therefore show derepression upon miRNA depletion even in *EZH2* -/- cells. Of the 213 genes co-repressed by *EZH2* and miRNAs, 116 genes (*P* = 8.17e-12) showed reduced derepression in response to miRNA depletion in *EZH2* -/- cells compared to WT cells (Fig. 4B-D), suggesting that PRC2 and miRNAs work coordinately, rather than independently, to repress these genes. Consistent with the heatmap (Fig. 4B), these feed-forward regulated genes showed derepression in response to miRNA depletion in WT but not in *EZH2* -/- cells (Fig. 4C). We confirmed this finding by qRT-PCR for several genes (Fig. 4E). The impaired derepression upon miRNA depletion in the absence of *EZH2* is also not due to genes attaining maximally derepressed expression levels in *EZH2* -/- cells. The remaining 47% of the genes co-repressed by *EZH2* and miRNAs that were not part of the feed-forward network showed significant derepression in response to miRNA loss both in WT and *EZH2* -/- cells, indicating that multiple levels of repression are detectable for genes that are not part of the coordinated feed-forward regulatory network (Supplemental Fig. S8).

**Figure 4.**
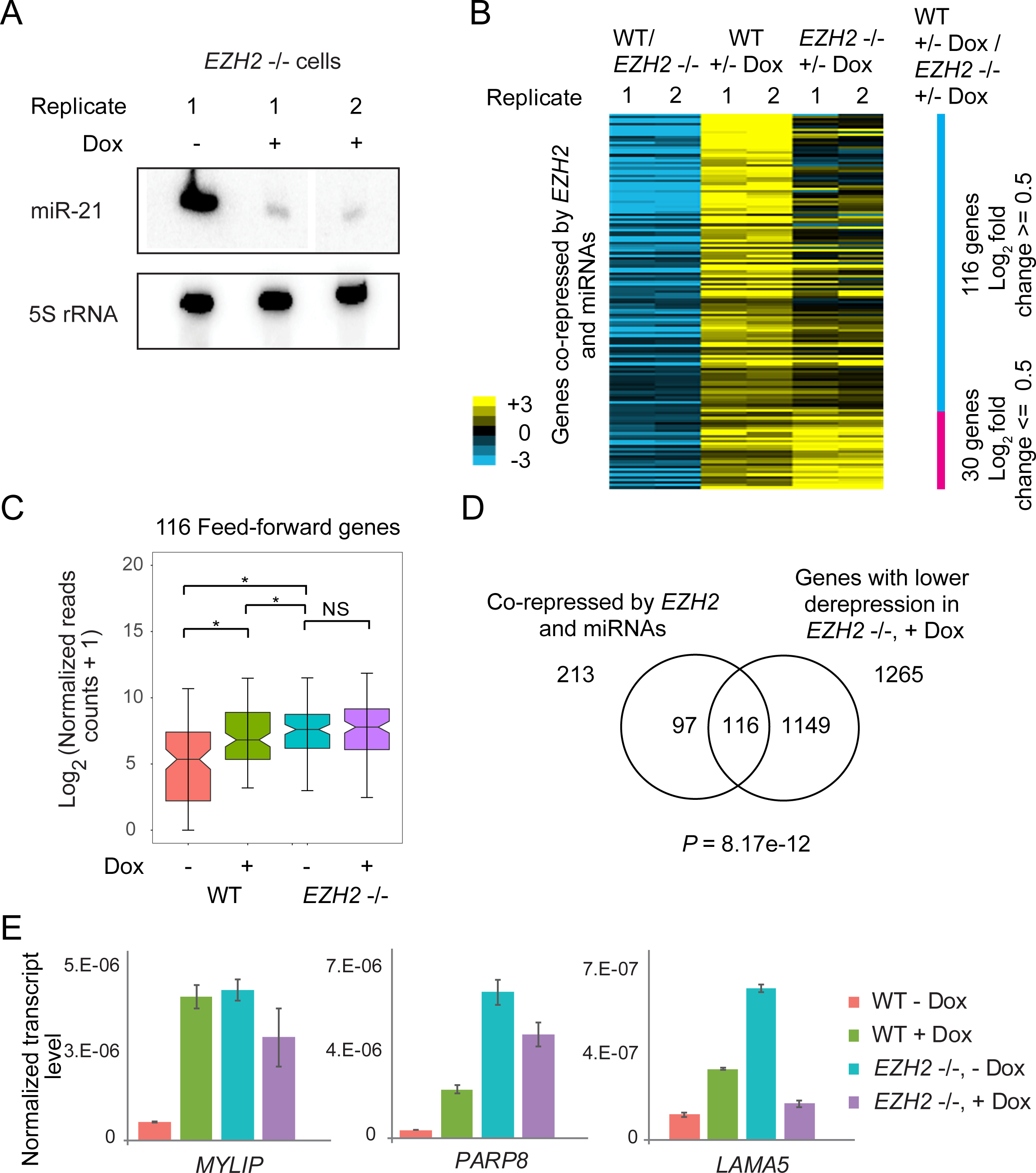
miRNAs repress PRC2 target genes through a feed-forward regulatory network with PRC2. (A) Depletion of miR-21 in response to *VP55* induction in *EZH2* -/- cells. (B) Heat map showing derepression of gene expression in response to miRNA loss (+/- Dox) in WT (WT, +/- Dox) and *EZH2* -/- cells (*EZH2* -/-, +/- Dox) for genes co-repressed by *EZH2* and miRNAs. Only 146 of the 213 co-repressed genes that had an average of 0.5 log_2_-fold difference in derepression between WT and *EZH2* -/- cells are shown. Genes are ordered in decreasing order of WT, +/- Dox/*EZH2* -/-, +/- Dox values. (C) Box plot comparing expression of feed-forward regulated genes in miRNA depleted (+ Dox) and non-depleted (- Dox) WT and *EZH2* -/- cells. * indicates *P* < 0.05 and NS indicates not significant. (D) Overlap between genes co-repressed by *EZH2* and miRNAs, and genes showing lower derepression upon miRNA depletion in *EZH2* -/- cells compared to WT cells. Significance of overlap was calculated using the hypergeometric test. (E) qRT-PCR showing transcript levels in miRNA-depleted WT and *EZH2* -/- cells, plotted as in Fig. 1E.

This coordination between PRC2 and miRNAs can be accounted for by a feed-forward regulatory network where *EZH2* promotes the miRNA-mediated repression of a subset (54%) of its direct targets, by activating miRNAs that repress those targets. Alternatively, it is possible that miRNAs somehow promote PRC2 function, perhaps by promoting the binding of PRC2 to its repression targets (Graham et al. 2016). To distinguish between these possibilities, we performed EZH2 ChIP-seq in cells overexpressing *VP55* (+ Dox) and untreated (-Dox) controls. EZH2 binding in the untreated cells (*VP55* integrated cell line) was similar to T98G WT cells (Supplemental Fig. S9A-C). If miRNAs promoted PRC2 binding at the feed-forward regulated genes, we would expect EZH2 binding to decrease in response to loss of miRNAs (+ Dox). However, there was no reduction in the level of EZH2 binding in Dox treated relative to untreated cells at the feed-forward regulated genes (Fig. 5A,B). We used qRT-PCR to verify derepression of key genes in response to miRNA depletion after Dox treatment in the same cultures of cells used for this ChIP-seq experiment (Supplemental Fig. S9D). These results confirm that miRNAs do not promote PRC2 binding or repression activity at the genes that are coordinately repressed by both miRNAs and PRC2, indicating instead that PRC2 promotes the miRNA-mediated repression of many of its targets.

**Figure 5.**
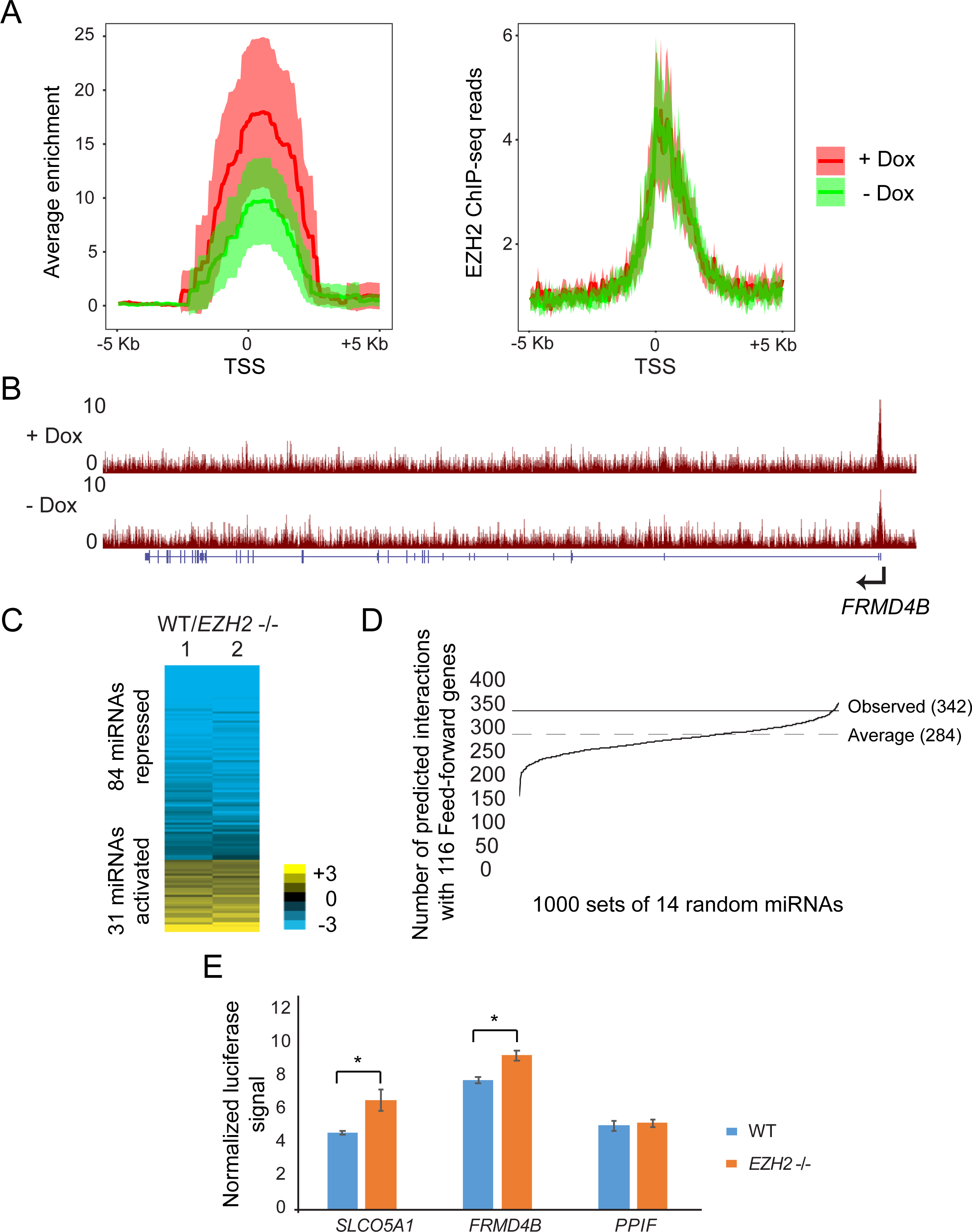
(A) Normalized density of EZH2 ChIP enrichment scores (-log_10_ *Q*-value calculated using MACS2, left) and EZH2 ChIP-seq reads (right) in the 10 kb region spanning the TSS of the feed-forward regulated genes in cells treated with Dox (+ Dox) and untreated (-Dox). (B) Genome browser view of *FRMD4B* showing tracks for EZH2 ChIP-seq in WT + Dox and WT - Dox cells containing *VP55*. (C) Heat map showing differential expression of miRNAs in response to *EZH2* knockout. Rows are ordered from low to high log_2_ fold change between WT and *EZH2* -/- miRNA counts. (D) Predicted interactions between 1000 random sets of 14 miRNAs and feed-forward regulated genes. The average across 1000 intersections is marked “Average”. The number of observed interactions between *EZH2*-activated miRNAs and feed-forward regulated genes is marked “Observed”. *P*-value = 0.024 was calculated using the bootstrap method (see methods). (E) Luciferase assay comparing miRNA inhibition of WT 3’ UTR transfected into WT and *EZH2* -/- cells.

Our data thus suggests a model where PRC2 transcriptionally represses hundreds of genes in GBM cells and for a significant fraction of these genes, it further promotes additional repression by activating miRNAs that post-transcriptionally repress those genes. To identify miRNAs regulated by PRC2, we performed miRNA-seq in WT and *EZH2* -/- cells and identified 31 miRNAs that were activated and 84 miRNAs that were repressed by *EZH2* (Fig. 5C). We found no EZH2 binding to chromatin around the *EZH2*-activated miRNAs, suggesting that *EZH2* activated them indirectly.

To determine whether the feed-forward regulated genes as a group were likely to be targeted by *EZH2*-activated miRNAs, we tested whether the number of predicted interactions between them were higher than expected by random chance. There were 342 unique interactions predicted between 14 *EZH2*-activated miRNAs (conserved miRNAs with mRNA target predictions) and 85 of the 116 feed-forward regulated genes. This observed number of interactions was significantly higher than expected by random chance, which we estimated using a bootstrapping approach (*P* = 0.024, see Methods, Fig. 5D). To verify that *EZH2* promoted post-transcriptional repression of its direct targets, we performed luciferase reporter assays in *EZH2* -/- and WT cells where we cloned the 3’ UTR spanning the AGO2 binding sites in two feed-forward regulated genes, *FRMD4B* and *SLCO5A1*. We detected lower luciferase activity in WT cells compared to *EZH2* -/- cells for both genes, pointing to higher post-transcriptional repression in the presence of *EZH2* (Fig. 5E). This implies that *EZH2* indirectly promotes expression of the miRNAs that are likely to post-transcriptionally regulate its direct targets.

### PRC2 regulates *FRMD4B* expression through a feed-forward regulatory network with let-7i

To verify the feed-forward regulatory network for a selected example in detail, we analyzed the repression of *FRMD4B* by PRC2. Consistent with direct transcriptional regulation by PRC2, both H3K27me3 and EZH2 were enriched near the promoter of *FRMD4B* (Fig. 6A). To verify that PRC2 promoted miRNA-mediated repression of *FRMD4B*, we measured its transcript levels in WT and *EZH2* -/- cells, with and without miRNA depletion. Loss of *EZH2* caused derepression of *FRMD4B*, whereas depletion of miRNAs caused derepression of *FRMD4B* expression in WT but not in *EZH2* -/- cells (Fig. 6B), verifying that PRC2 promotes miRNA-mediated regulation of *FRMD4B*. To test if PRC2 promotes expression of a specific miRNA that targets *FRMD4B*, we first identified let-7i as an *EZH2*-activated miRNA with a predicted interaction site near an AGO2 iCLIP site on the *FRMD4B* transcript (Fig. 6C and Supplemental Fig. S10A). We then analyzed the regulation of *FRMD4B* by let-7i using luciferase assays with a WT or mutant *FRMD4B* 3’ UTR in WT and *EZH2* -/- cells. Deletion of the let-7i binding site led to a marked increase in luciferase signal in WT but a much smaller increase in *EZH2* -/- cells (*P* = 0.0098, paired t-test, Fig. 6D). Thus, *FRMD4B* is more strongly repressed by let-7i in the presence of its activator EZH2.

**Figure 6.**
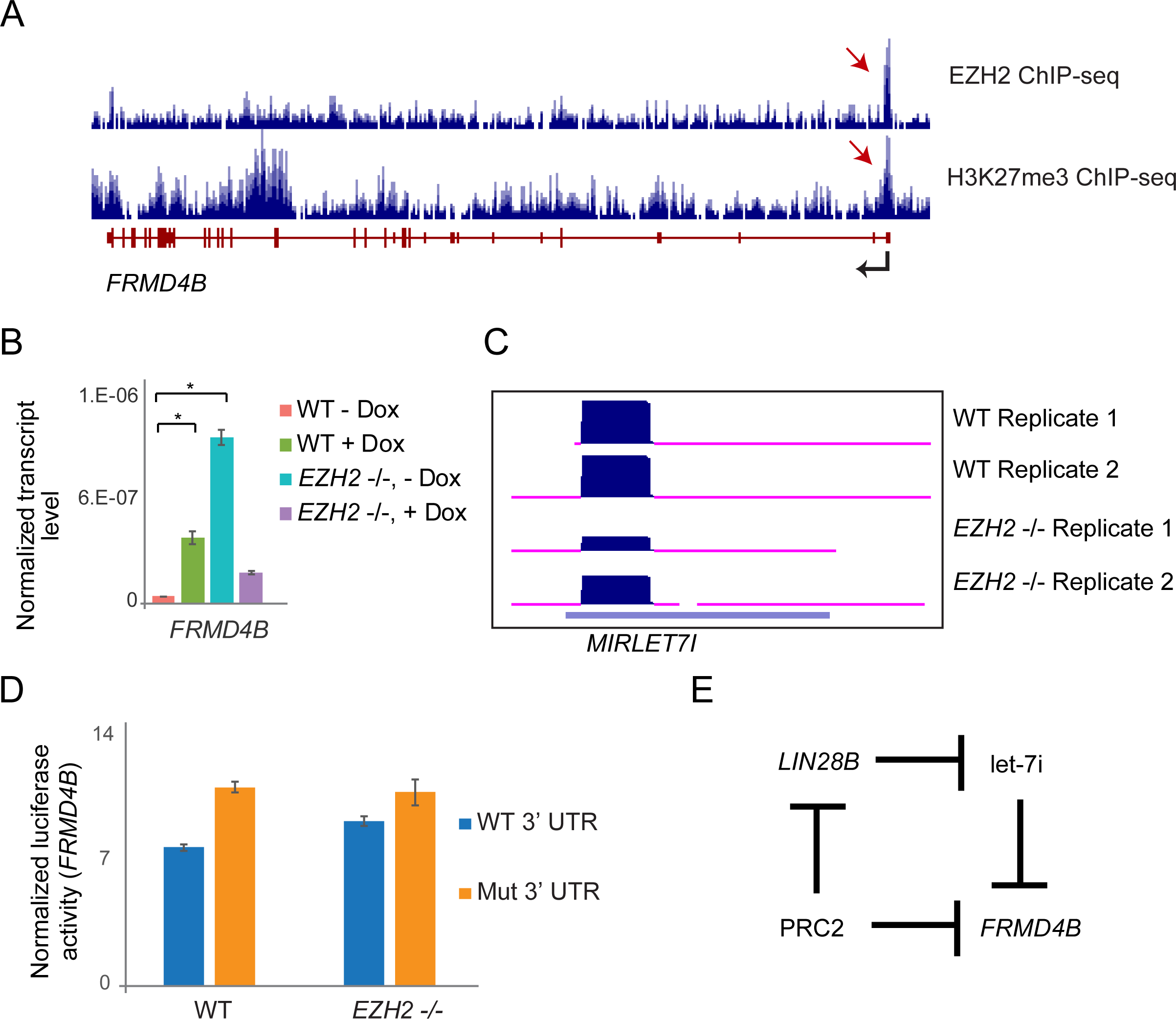
PRC2 regulates *FRMD4B* expression through a feed-forward regulatory network with let-7i. (A) Genome browser view of *FRMD4B* showing tracks for EZH2 and H3K27me3 ChIP-seq. (B) qRT-PCR showing expression changes of *FRMD4B* in miRNA-depleted (+ Dox) and non-depleted (- Dox) WT and *EZH2* -/- cells, plotted as in Fig. 1E. * indicates *P* < 0.05. (C) Genome browser view of let-7i in two replicates of WT and *EZH2* -/- cells. (D) Luciferase assay comparing miRNA inhibition between WT and mutant *FRMD4B* 3’ UTR (seed deletion) transfected WT and *EZH2* -/- cells. (E) Feed-forward regulatory network formed by PRC2, *LIN28B*, let-7i and *FRMD4B*.

To address how *EZH2* indirectly activates let-7i, we searched for likely repressors of let-7i among the genes directly repressed by *EZH2*. We found that *LIN28B*, a known repressor of the let-7 family of miRNAs, showed chromatin marks characteristic of *EZH2* activity (Supplemental Fig. S10B). Our RNA-seq data showed that *LIN28B* was significantly derepressed upon loss of *EZH2* (FDR corrected *P* < 0.05), suggesting that it was directly repressed by *EZH2*. LIN28B is an RNA-binding protein that inhibits the biogenesis of let-7 miRNAs (Balzeau et al. 2017). Since *LIN28B* is known to be a common post-transcriptional repressor of the let-7 family, *EZH2* repression of *LIN28B* might be expected to activate multiple members of the let-7 family. Indeed, several other let-7 miRNAs also showed derepression in *EZH2* -/- cells, though only let-7i and let-7f-3p met the stringent threshold for statistical significance we used earlier to identify derepressed miRNAs (Supplemental Fig. S10C). These data suggest that *EZH2* activates let-7i and other let-7 miRNAs by directly repressing *LIN28B* (Fig. 6E). Taken together, these results confirm that PRC2 represses *FRMD4B* expression through a feed-forward regulatory network with let-7i.

### Discussion

Transcription factors and miRNAs work in a highly coordinated manner, forming feed-back and feed-forward networks to regulate diverse processes. Using a combination of crosslinking based immunoprecipitation (iCLIP) and global miRNA depletion followed by RNA-seq, we identified miRNA-regulated genes genome-wide in GBM cells. This approach however only identified genes whose transcript levels are lowered due to miRNA function and thus likely underestimates the number of miRNA-repressed genes. Hundreds of miRNA-repressed genes nonetheless showed enrichment for PRC2 binding and H3K27me3 marks. We found that miRNAs post-transcriptionally reinforce the transcriptional repression of a significant fraction of PRC2 target genes, either independently of PRC2, or coordinately, by forming a feed-forward regulatory network with PRC2.

Overall, genes repressed by miRNAs and by PRC2 constituted 22% of all direct PRC2-repressed genes. 54% of these genes (feed-forward regulated genes) showed much less derepression in response to miRNA depletion in the absence of *EZH2*, indicating that PRC2 contributed to the repression function of some of these miRNAs. Although our data suggests miRNA-mediated post-transcriptional repression of PRC2 direct targets, we could verify AGO2 binding by iCLIP for only 34.7% of the genes co-repressed by *EZH2* and miRNAs (Supplemental Fig. S6B). This can be attributed to miRNA-independent binding of AGO2 and/or to technical limitations of iCLIP in capturing low-abundance transcripts (Leung et al. 2011; Müller-McNicoll et al. 2016; Bieniasz and Kutluay 2018). The difference in derepression in *EZH2* -/- cells was not due to reduced miRNA depletion by *VP55* in *EZH2* -/- cells since the extent and the number of miRNAs depleted in WT and *EZH2* -/- cells were almost identical (Supplemental Fig. S7B,C). The coordinated feed-forward repression by miRNAs and PRC2 was not due to a mechanism in which miRNAs promote PRC2 function or binding (Graham et al. 2016), because loss of miRNAs had no effect on PRC2 binding at the feed-forward regulated genes (Fig. 5A,B).

We propose that many genes that are transcriptionally repressed by PRC2 continue to produce transcripts at low levels due to inefficient repression by PRC2 at the chromatin and transcriptional level. These transcripts are then further repressed or destabilized by miRNAs which thus reinforce the repressive function of PRC2 in one of two modes. In the coordinated mode, miRNAs that repress PRC2 targets are themselves indirectly activated by *EZH2*, forming feed-forward regulatory networks. In the independent mode, miRNAs and PRC2 co-repress a common set of target genes independently (Fig. 7). This model would be in agreement with the notion that miRNAs are broadly involved in reinforcing transcriptional programs (Ebert and Sharp 2012). We found that PRC2 binds and tri-methylates H3K27 to repress genes that also harbor active H3K4me3 marks. These genes could produce residual transcripts as a result of leaky transcription, and indeed we found this to be the case. Genes co-repressed by PRC2 and miRNAs showed significantly higher expression upon depletion of miRNAs than genes silenced by PRC2 alone, even though PRC2 showed equal occupancy at both sets of genes (Supplemental Fig. S11).

**Figure 7.**
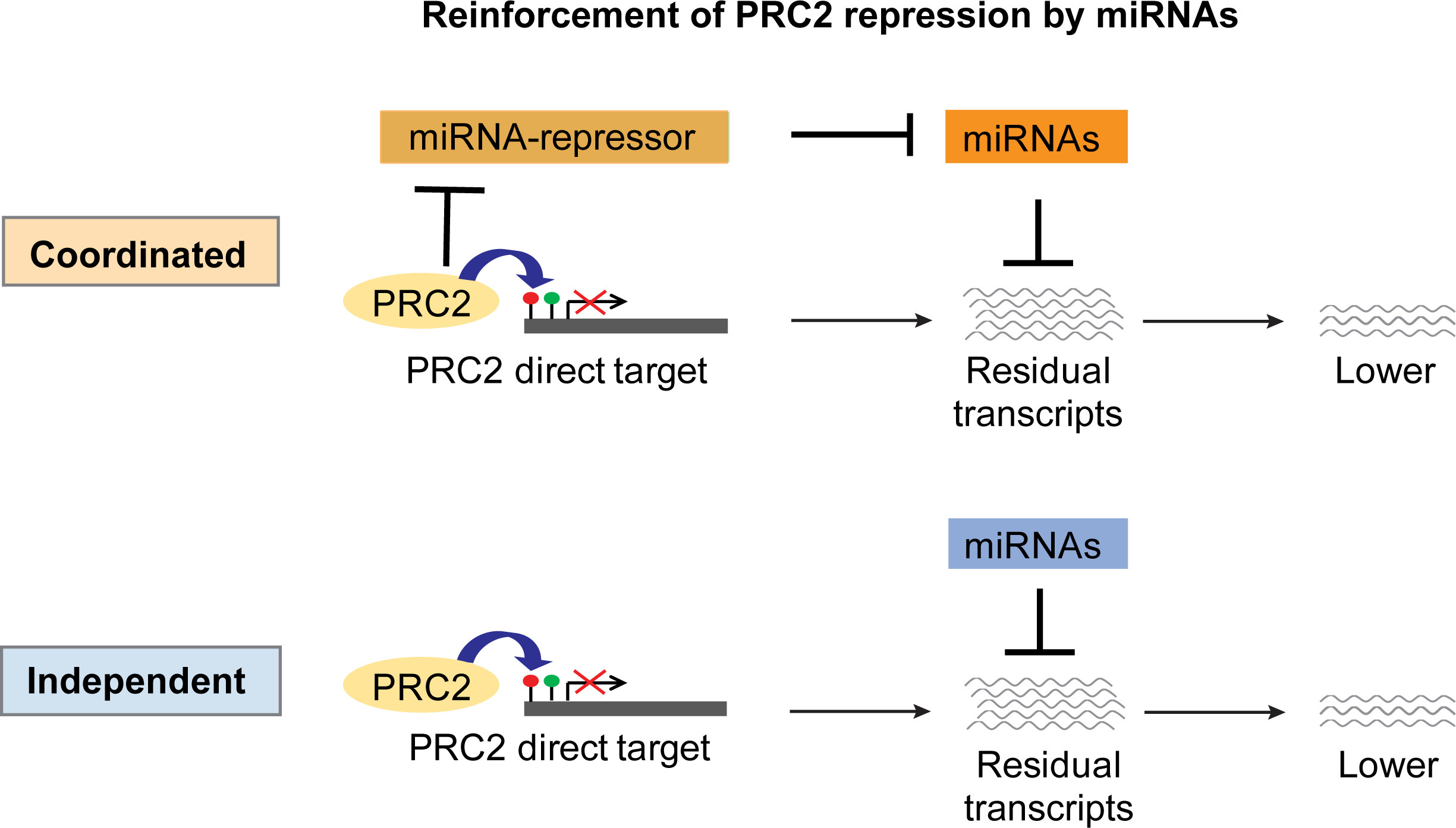
Model for reinforcement of PRC2 repression by miRNAs. Co-repression of target genes occurs transcriptionally by PRC2 and post-transcriptionally by miRNAs, either in a coordinated or independent manner.

It is likely that this coordination between PRC2 and miRNA repression occurs in other cellular contexts, given the significant overlap in the genes regulated by PRC2 and miRNAs in mESCs (Supplemental Fig. S4). Our findings not only uncover a novel type of coordination between an epigenetic and post-transcriptional regulator of gene expression but could also have a broad impact on PRC2’s regulatory function in a wide range of disease and developmental pathways.

## Methods

### Cell lines and reagents

GBM cell lines T98G and U87MG (ATCC-CRL-1690 and ATCC-HTB14) were grown in EMEM with 10% FBS (Gibco). All cell lines were maintained at 37oC and 5% CO_2_. Antibodies used were – EZH2 (Abcam 186006), AGO2 (Abcam, ab 57113), SUZ12 (Active motif 39877), H3K27me3 (Millipore 07-449) and PPIF (Abcam, ab 110324). pCMV(CAT)T7-SB100 was a gift from Zsuzsanna Izsvak (Addgene plasmid # 34879). pSBtet-GP was a gift from Eric Kowarz (Addgene plasmid # 60495).

### RNA-seq, miRNA-seq and iCLIP

RNA-seq experiments were performed on poly(A) selected mRNA (Bioo 512980) as previously described (Hall et al. 2018). AGO2 iCLIP experiments were performed as previously described (König et al. 2010). Small RNA-seq libraries were prepared using NEBNext small RNA library preparation kit (NEB E7330) as previously described (Shivram and Iyer 2018).

### Quantitative Reverse Transcription PCR (qRT-PCR)

Total RNA was extracted using TRIzol and reverse transcribed by Superscript III reverse transcriptase (Thermo Fisher Scientific 18080085) using random hexamer primers. PCR was then performed on cDNA using Power SYBR Green master mix (Thermo Fisher Scientific 4367659). Transcript levels were normalized to 18S RNA levels and fold changes were quantified using the ΔΔCt method (Livak and Schmittgen 2001).

### Chromatin immunoprecipitations

Cells were crosslinked with 1% formaldehyde for 20 min and quenched with 125 mM glycine. The rest of the protocol was performed as previously described (Hall et al. 2018).

### mRNA-seq and miRNA-seq analysis

Reads were aligned to the human genome (UCSC version hg38) using Hisat (Kim et al. 2015). FeatureCounts was then used to count reads mapping to genes and differential gene expression analysis was performed using DESeq2 with default parameters (Liao et al. 2014). Significantly differentially expressed genes were identified at a threshold of FDR corrected *P* < 0.05 and log_2_ fold-change ≥ 0.5 or ≤ −0.5. For miRNA-seq, reads mapping to mature miRNAs (miRBase hg38) were counted using BEDTools (Quinlan and Hall 2010). Differentially expressed miRNAs were then identified using DESeq2 as described above, using the same thresholds (Love et al. 2014). Feed-forward target genes are those that show at least log_2_-fold 0.5 higher miRNA-mediated repression in T98G WT cells compared to *EZH2* -/-.

### ChIP-seq analysis

ChIP-seq reads were aligned to the human genome (UCSC version hg38) using BWA (Li and Durbin 2009). MACS2 was then used to call peaks at the cut-off of *P*-value < 0.01 (Zhang et al. 2008). Genes containing an EZH2 peak within 10 kb around the TSS were used as EZH2-bound genes for downstream analysis. For the heat maps plotted in Fig. 3B, Fig. 3D and Supplemental Fig. S5B, the signal represents enrichment scores (-log_10_ *Q*-value) output by MACS2. ChIP-seq data for H3K4me3 in GBM cells was obtained from our recently published study (Hall et al. 2018).

### iCLIP analysis

iCLIP reads were first processed to remove duplicate reads using FASTX Toolkit (http://hannonlab.cshl.edu/fastx_toolkit/) and then aligned to the human genome (UCSC version 19) using Bowtie 2. Reads mapping to repeat regions were filtered out using RepeatMasker (Smit et al. 2013–2015). Peaks were then called using CLIPper using default parameters and filtered based on reproducibility across replicates (Lovci et al. 2013). MiRNA seed enrichments in sequences spanning 200 bases around peaks were determined using HOMER software (Heinz et al. 2010). Prior to intersections with other datasets from RNA-seq, peak coordinates were converted to hg38 from hg19 using liftOver.

### CRISPR-Cas9 knockout of *EZH2*

*EZH2* sgRNA (sequence TTATCAGAAGGAAATTTCCG) was designed using http://crispr.mit.edu, and *EZH2* -/- T98G cells were generated using the GeneArt CRISPR-Cas kit (Thermo Fisher Scientific A21175) following manufacturer’s instructions. Knockout clones were verified by immunoblots.

### Generation of *VP55*-expressing stable cell line

The *VP55* coding sequence was PCR-amplified from a vector containing the codon-optimized *VP55* sequence (a gift from Christopher Sullivan and Benjamin tenOever) and cloned into pSBtet-GP, a Tetracycline/Doxycycline-inducible expression vector containing the *Sleeping Beauty* transposase-specific inverted terminal repeats flanking the cloning site. Introduction of pSBtet-GP-*VP55* and the *Sleeping Beauty* transposase expressing vector (SB100X) into cells allows the transposase-mediated genomic integration of the DNA sequence from pSBtet-GP-*VP55* that is flanked by the inverted terminal repeats. SB100X and pSBtet-GP-*VP55* constructs were transfected into T98G cells using Lipofectamine 2000 as per manufacturer’s instructions (Thermo Fisher Scientific 11668030). Clonal cells were then selected through puromycin selection at 2.5 µg/ml (Gibco A1113803). Expression of *VP55* was induced with Doxycycline (D3072) at 5 µg/ml.

### Luciferase reporter assays

The 3’ UTR of target mRNAs spanning at least 500 bp around the predicted binding site of selected miRNAs were cloned into the psi-CHECK2 vector downstream of the Renilla luciferase gene. Mutagenesis was performed using the QuikChange II site directed Mutagenesis Kit (Agilent, 200523). Cells were transfected with WT or mutant 3’ UTR with or without miRNA inhibitors and harvested after 48 hours. Luciferase activity was then measured using the Promega dual luciferase kit and calculated as the ratio of luminescence from Renilla to Firefly.

### miRNA target enrichment analysis

1000 random sets of 14 miRNAs were generated from among miRNAs expressed in T98G cells. For each set, the number of predicted interactions with 116 feed-forward related genes were determined using the DIANA miRNA-mRNA interaction annotations (Paraskevopoulou et al. 2013). *P*-value was estimated from the fraction of iterations with ≥ 342 interactions that were observed for *EZH2*-activated miRNAs.

### Gene set overlaps using bootstrapping

10,000 random sets of 1834 or 959 genes were generated from a pool of genes not repressed by *EZH2* but expressed at similar levels as the *EZH2*-repressed genes. Overlaps were then calculated between each of the 10,000 sets and 3844 miRNA-repressed genes. *P*-value was estimated from the fraction of iterations with ≥ 444 genes.

### Data Access

Primary sequencing data generated in this study have been submitted to the NCBI Gene Expression Omnibus (GEO; http://www.ncbi.nlm.nih.gov/geo/) under accession number GSE112242. Scripts used in this manuscript are available at Github (https://github.com/haridh/PRC2-miRNA-manuscript).

## Acknowledgments

We thank Anna Battenhouse for assistance with aligning sequencing data and ChIP-seq analysis, Alan Lambowitz for the use of their UV254 crosslinker, Benjamin tenOever for the *VP55* construct, Chris Sullivan for suggesting the use of *VP55*, providing constructs and for discussions, Luiz Penalva and Suzanne Burns for advice on the iCLIP experiments, and the Genomic Sequencing and Analysis Facility at UT Austin and the MD Anderson Cancer Center-Science Park NGS Facility for Illumina sequencing. The Science Park NGS Facility was supported by CPRIT Core Facility Support Grants RP120348 and RP170002. We also thank the Texas Advanced Computing Center (TACC) at UT Austin for the use of computational facilities. This work was funded in part by grants from the Cancer Prevention and Research Institute of Texas (RP120194) and the National Institutes of Health (NIH) (CA198648) to V.R.I.

## Supplemental Material

Supplemental Figures S1-S11.

---

**Supplemental Figure S1.**
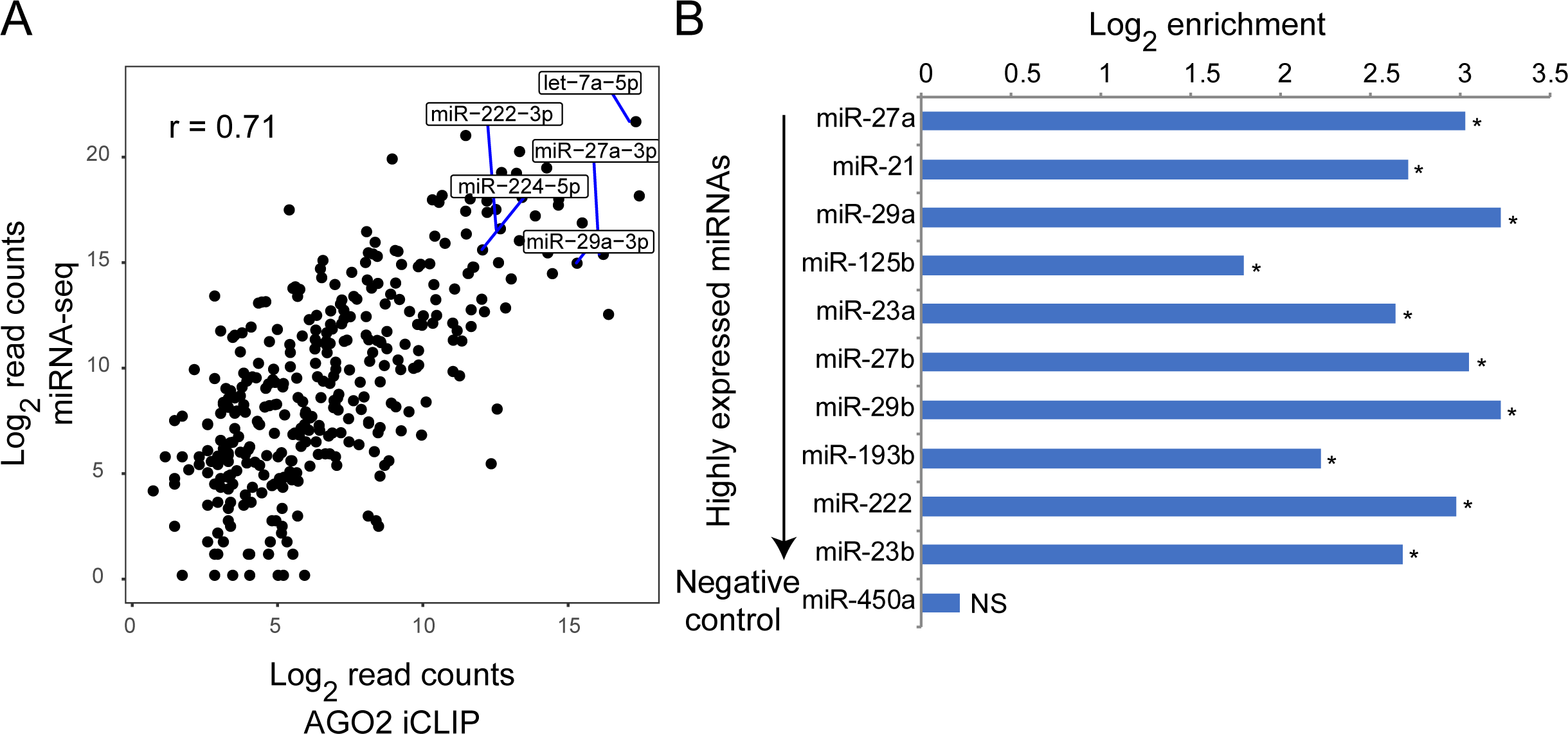
AGO2 binding sites identified by iCLIP are enriched for binding sites of highly expressed miRNAs. (A) Correlation of miRNA enrichment in AGO2 iCLIP data (X-axis) with their expression (Y-axis). AGO2 iCLIP-enriched miRNAs shown in Fig 1D (right) are indicated. (B) Enrichment for predicted mRNA targets of AGO2-interacting miRNAs among protein-coding genes interacting with AGO2.

**Supplemental Figure S2.**
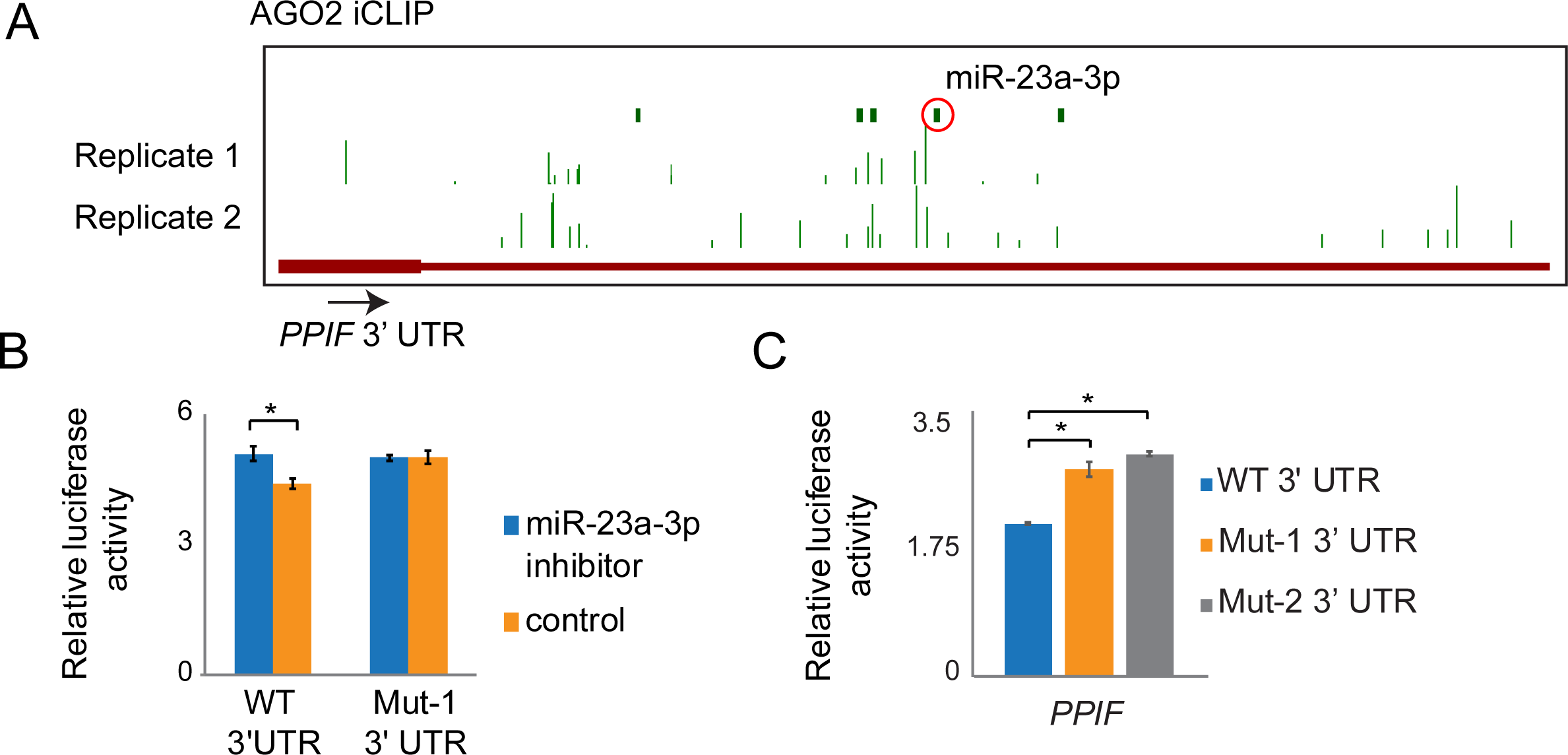
miR-23a-3p directly represses *PPIF* expression. (A) Genome browser view showing AGO2 iCLIP peaks at the 3’ UTR of *PPIF*. Green bars represent predicted miRNA binding sites with miR-23a-3p binding site highlighted in red. (B) Luciferase assays showing rescue of miRNA inhibition on treatment with miR-23a-3p inhibitor or negative control inhibitor for WT 3’ UTR and mutant 3’ UTR (seed deletion, Mut-1) transfected U87MG cells. (C) Changes in luciferase signal on deletion of miR-23a-3p binding site (Mut-1) or disruption of miRNA-*PPIF* interactions through base substitutions (Mut-2) in T98G cells.

**Supplemental Figure S3.**
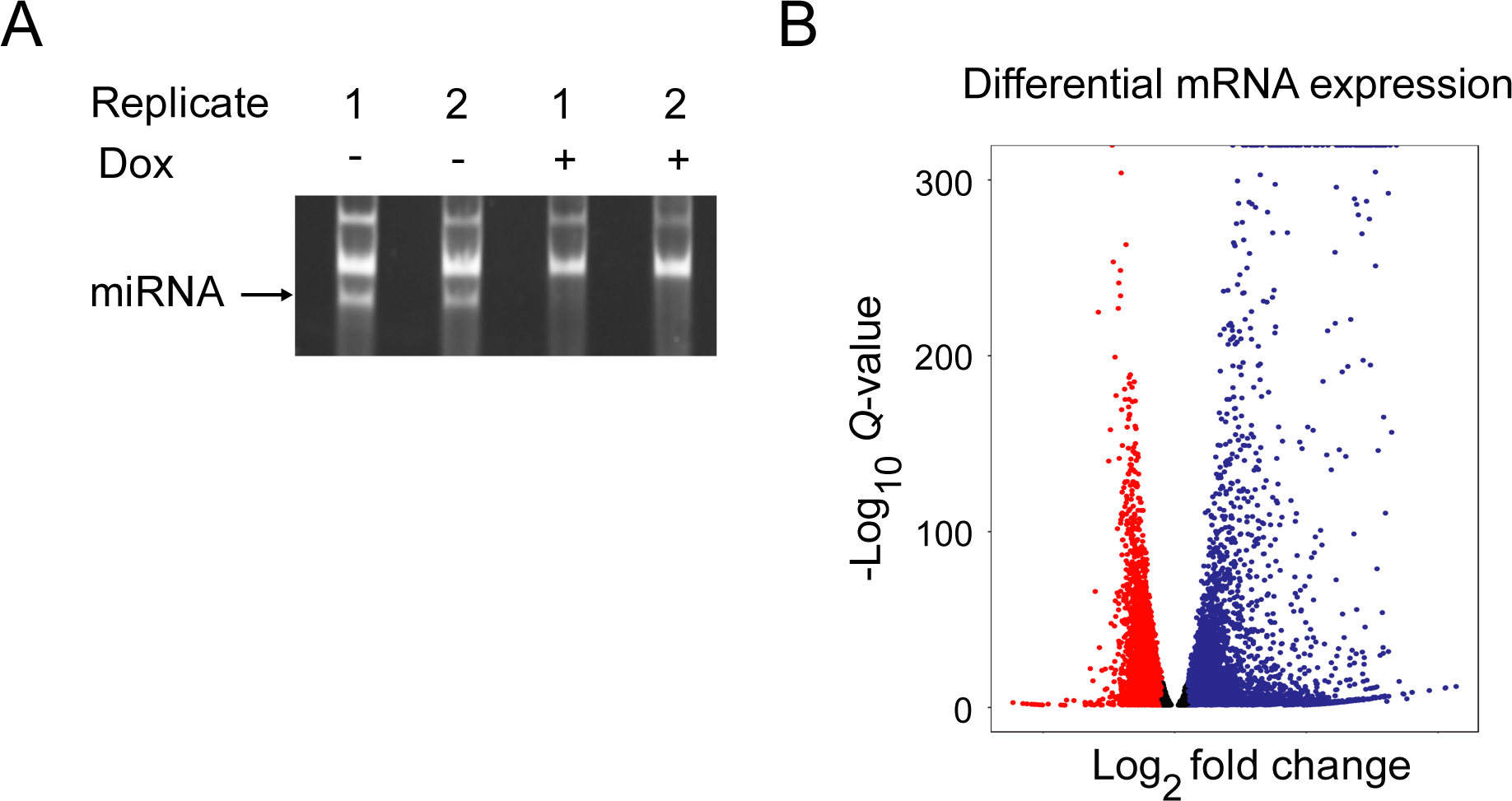
Identification of miRNA-repressed genes by global depletion of miRNAs. (A) cDNA libraries prepared from miRNA depleted (+ Dox) and non-depleted (-Dox) T98G cells. (B) Volcano plot of differentially expressed protein-coding genes in response to VP55 induction.

**Supplemental Figure S4.**
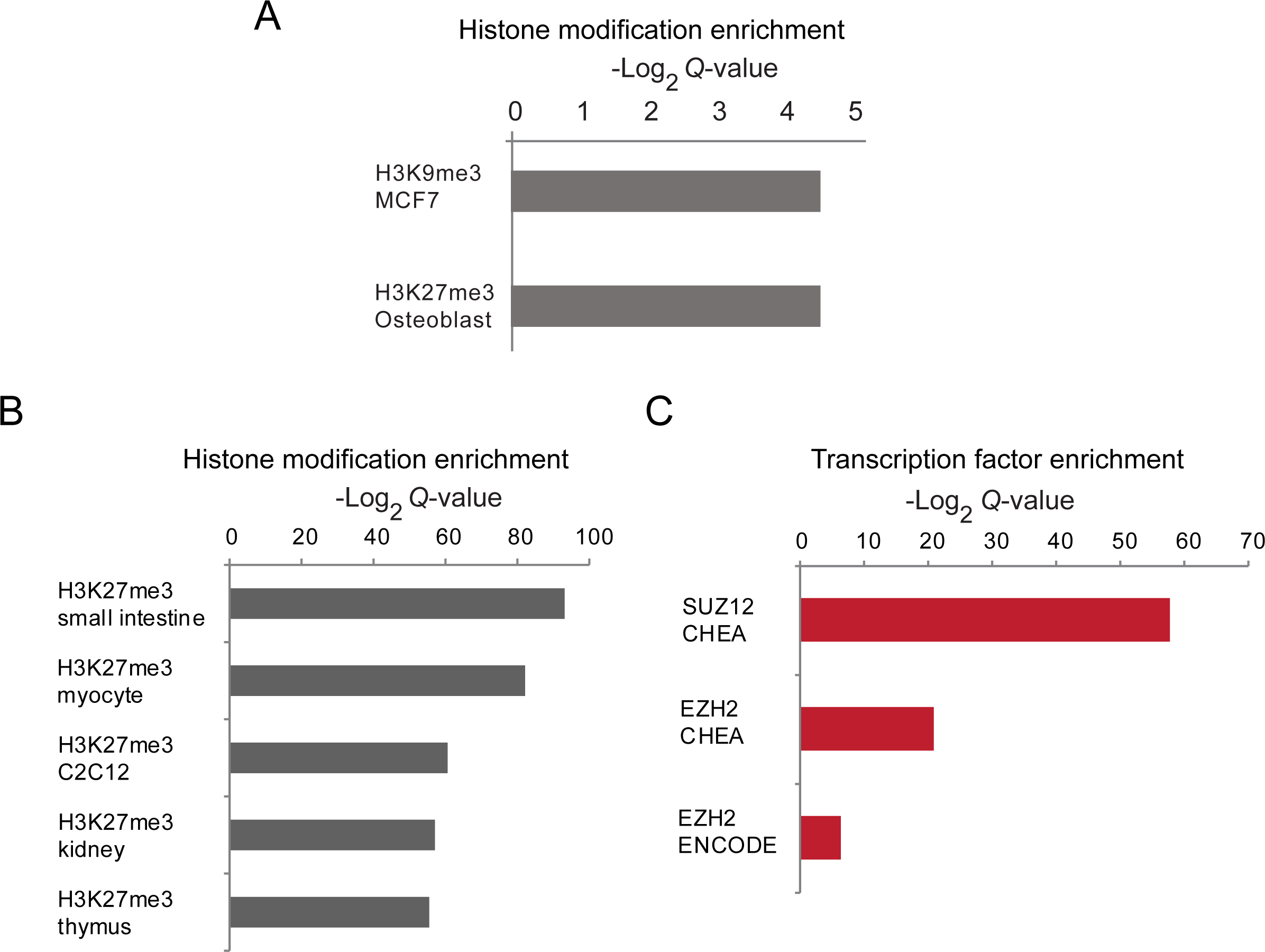
Gene set enrichment analysis using Enrichr. (A) Enrichr analysis showing enrichment for histone modifications at a control set of genes expressed in the same range as the miRNA-repressed genes identified in T98G cells. (B,C) Enrichr analysis showing enrichment for H3K27me3 (B) and PRC2 (C) at genes up-regulated in miRNA-depleted mESCs (DICER knockout mESCs).

**Supplemental Figure S5.**
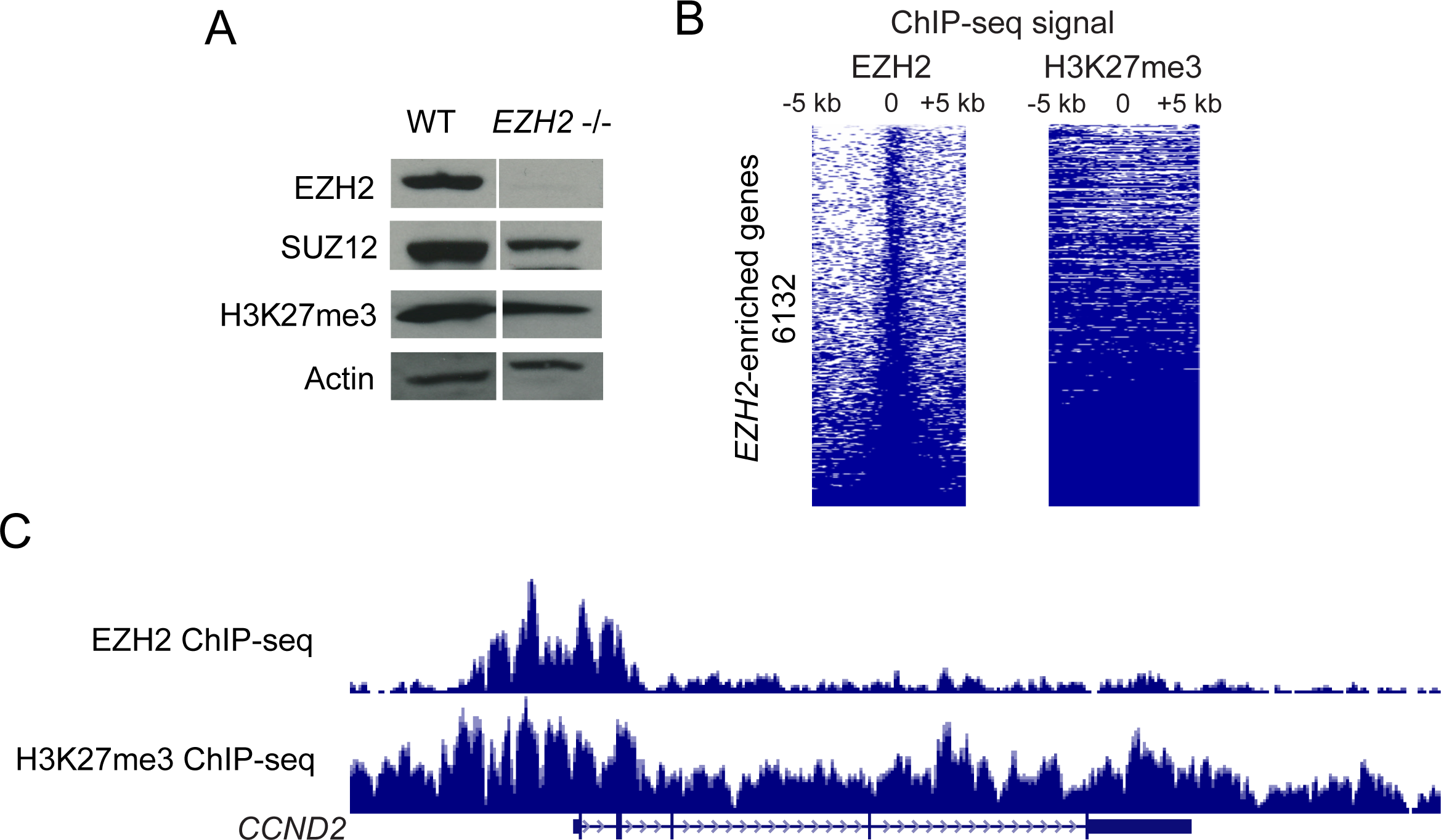
(A) Immunoblots for EZH2, SUZ12 and H3K27me3 in WT and *EZH2* -/- cells. (B) EZH2 and H3K27me3 ChIP-seq enrichment scores for protein-coding genes containing at least one significant EZH2 ChIP peak. Each row represents a gene containing at least one EZH2 ChIP-seq peak and the columns are binned into 50 bp windows spanning 10 kb around the TSS. (C) Genome browser view of *CCND2*, a gene enriched with EZH2 and H3K27me3 on chromatin, and that is derepressed in *EZH2* -/- cells.

**Supplemental Figure S6.**
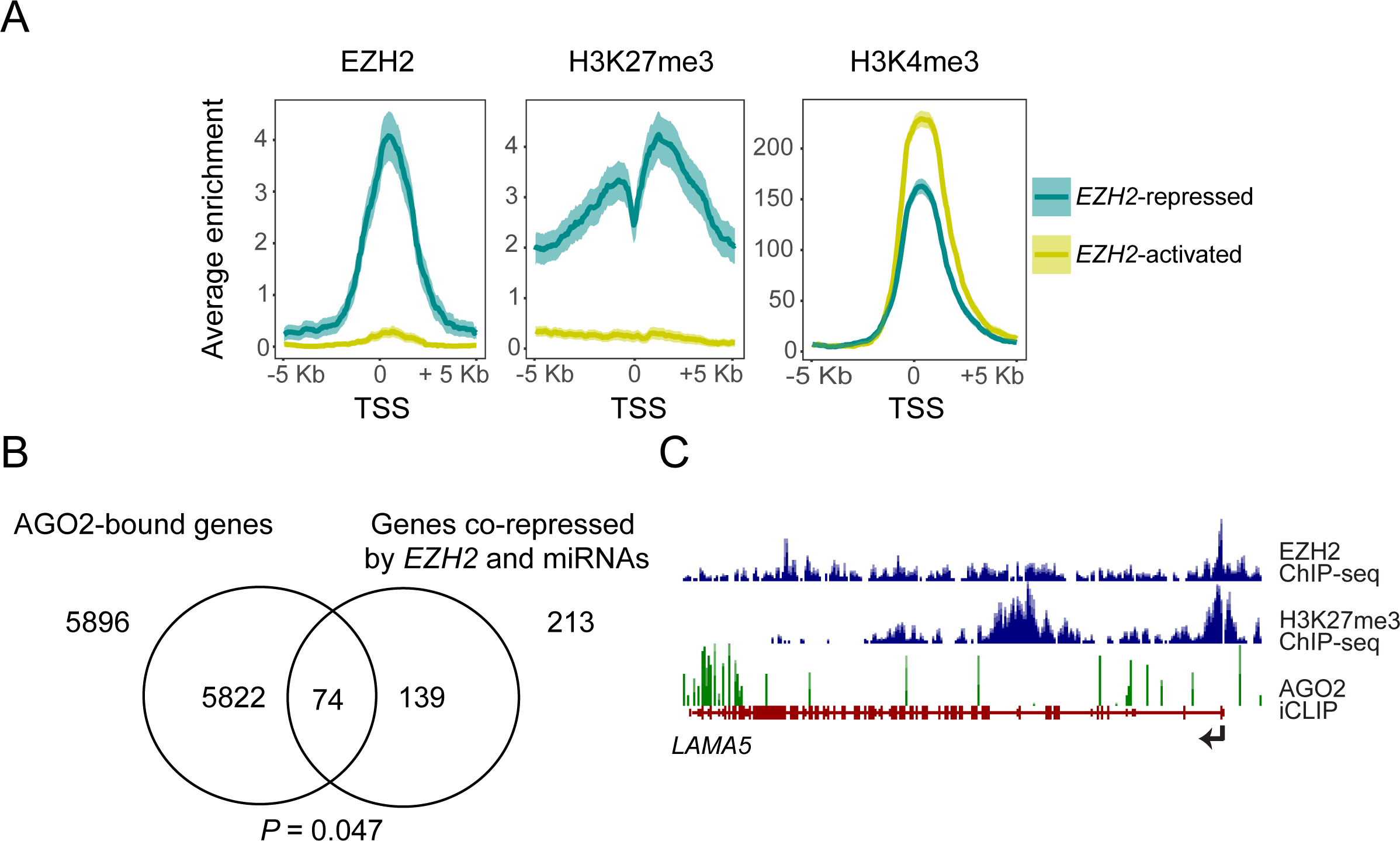
Genes derepressed on loss of EZH2 are enriched for PRC2 and also regulated by miRNAs. (A) Normalized density of EZH2, H3K27me3 and H3K4me3 ChIP enrichment scores in the 10 kb region spanning the TSS of EZH2-regulated genes plotted in Fig. 3A. (B) Overlap of AGO2-interacting genes repressed by miRNAs and genes co-repressed by *EZH2* and miRNAs. Significance of overlap was calculated using the hypergeometric test.(C) Genome browser view of *LAMA5*, a gene enriched with EZH2 and H3K27me3 on chromatin, and whose transcript is also bound by AGO2.

**Supplemental Figure S7.**
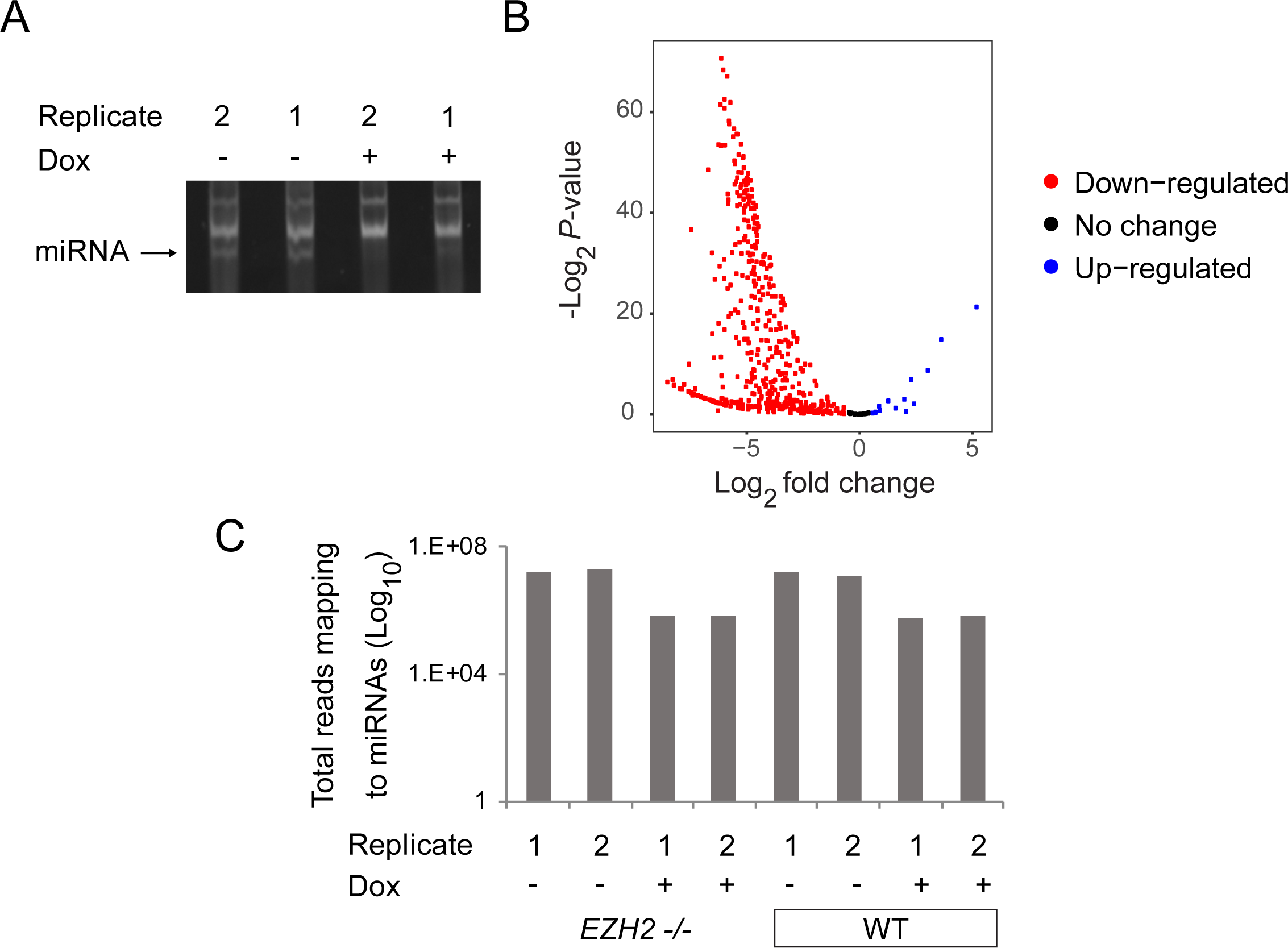
Global miRNA depletion in *EZH2* -/- cells is comparable to WT cells. (A) cDNA libraries prepared from *EZH2* -/- cells with (+ Dox) and without depletion of miRNAs (- Dox). (B) Volcano plot of differentially expressed miRNAs in response to *VP55* induction. X-axis represents log2 fold difference between expression of miRNAs in *VP55*-induced and non-induced cells. Y-axis represents −log2 P-value of the expression difference calculated using DESeq2. (C) Total miRNA counts (Log_10_ total reads mapping to miRNAs) in WT and *EZH2* -/- cells with and without depletion of miRNAs.

**Supplemental Figure S8.**
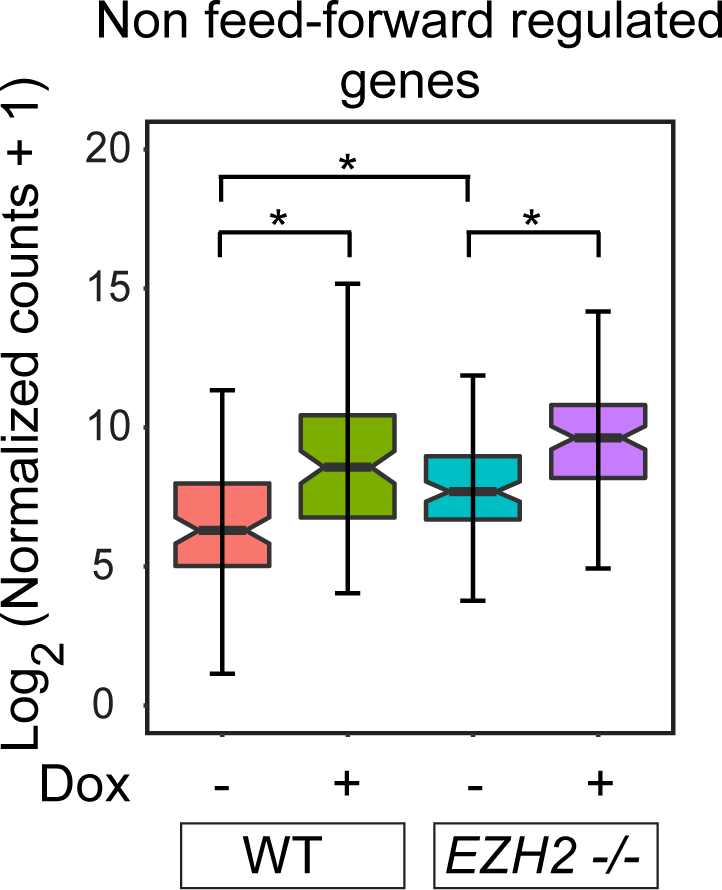
Non feed-forward regulated genes show derepression in response to miRNA loss in *EZH2* -/- cells and WT T98G cells. Box plot comparing expression of non-feed-forward regulated genes in miRNA depleted and non-depleted WT and *EZH2* -/- cells.

**Supplemental Figure S9.**
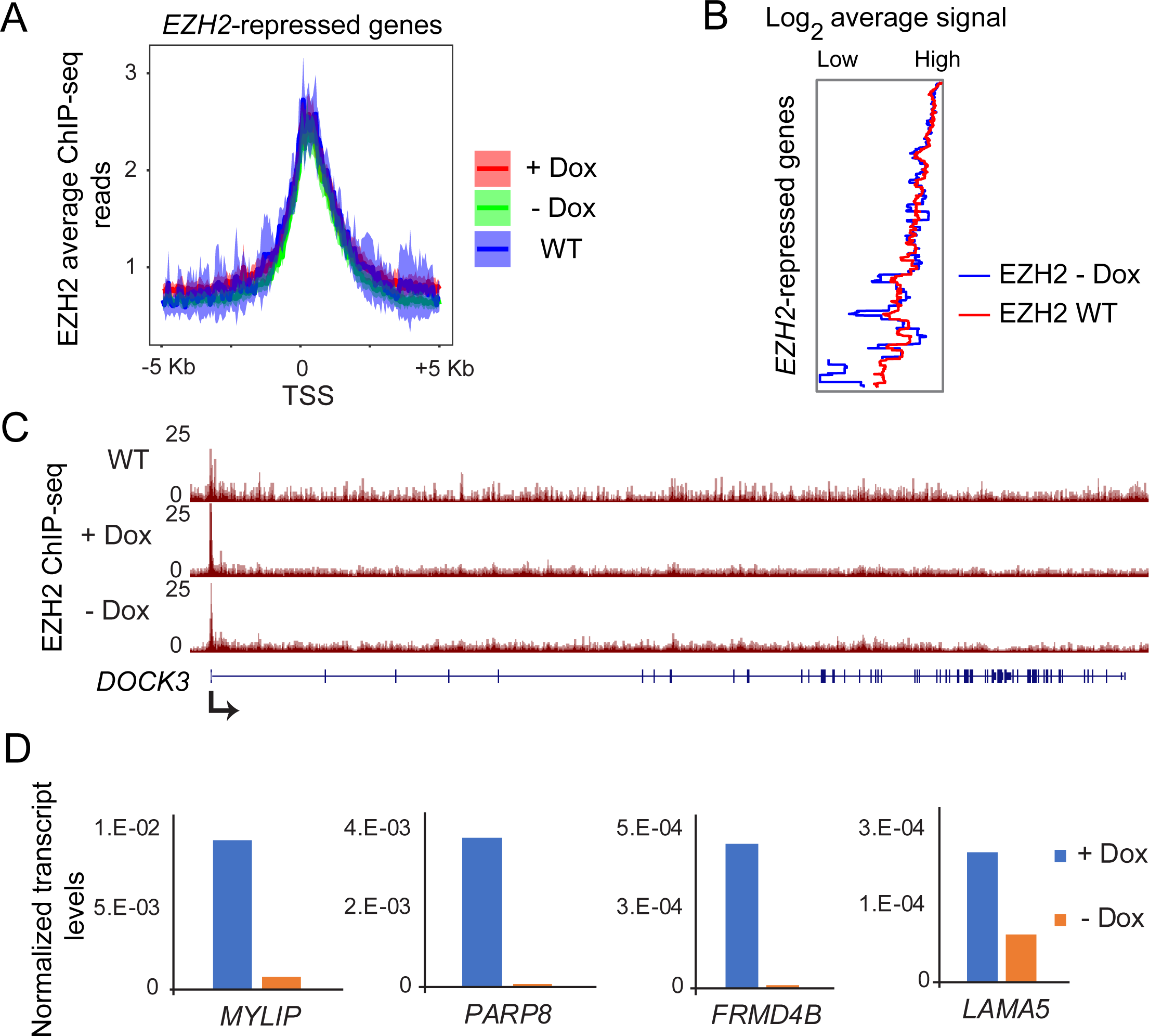
EZH2 ChIP-seq in *VP55*-integrated WT - Dox cells is consistent with EZH2 ChIP-seq in WT cells. (A) Normalized density of EZH2 ChIP-seq reads at *EZH2*-repressed genes from EZH2 ChIP-seq in *VP55*-integrated WT + Dox, WT - Dox and WT cells. (B) Average EZH2 ChIP-seq signal for genes plotted in B. (C) Genome browser view of *DOCK3*, showing EZH2 enrichment in WT T98G cells, WT + Dox and WT - Dox cells. (D) qRT-PCR validation of Dox-induced derepression of miRNA-repressed genes, plotted as in Fig. 1E and normalized to Actin.

**Supplemental Figure S10.**
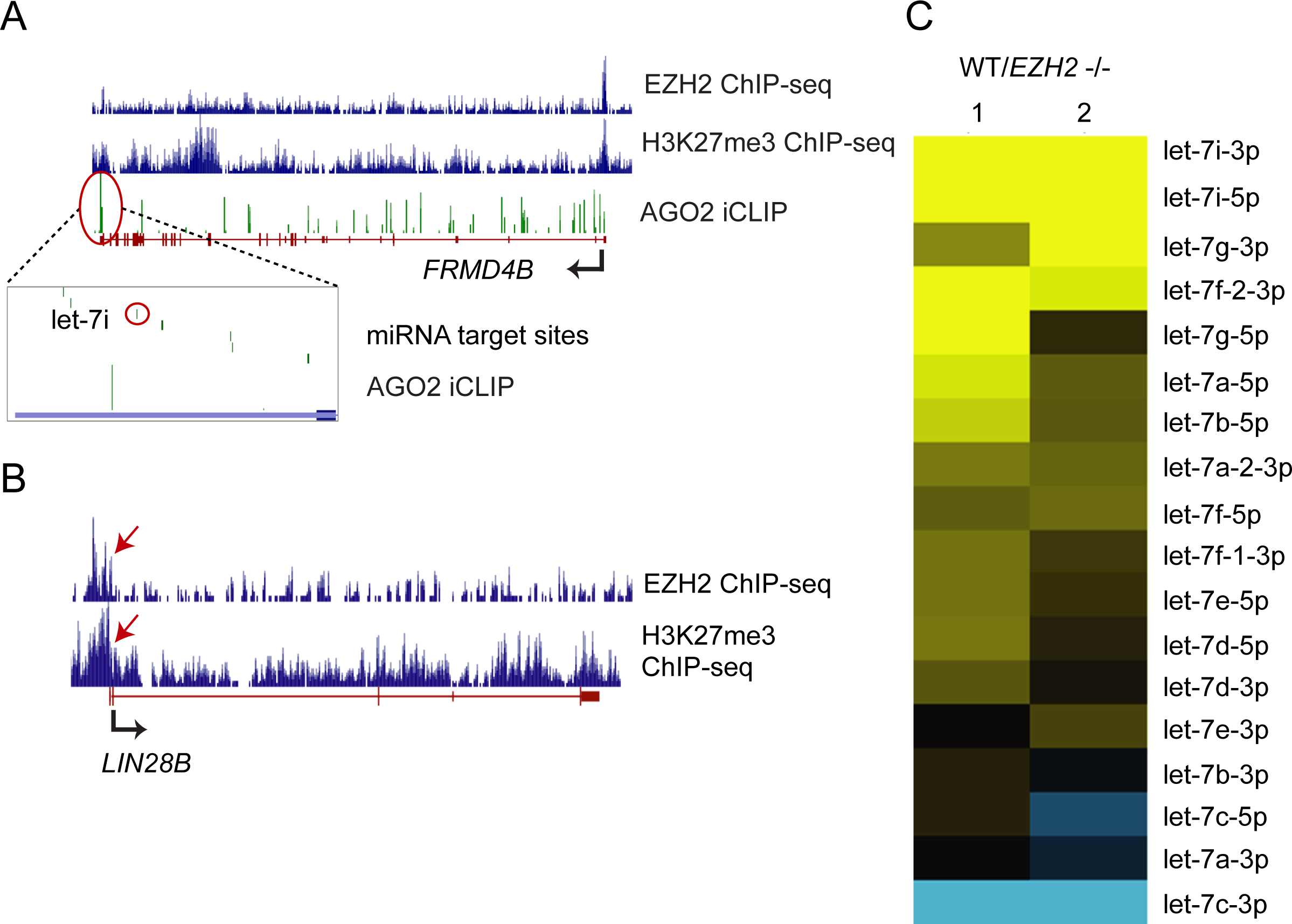
*EZH2* activates let-7 miRNAs by directly repressing *LIN28B*. (A) Genome browser view of *FRMD4B* showing tracks for H3K27me3 ChIP-seq, EZH2 ChIP-seq and AGO2 iCLIP highlighting the binding site for let-7i. (B) Genome browser view of *LIN28B* showing tracks for EZH2 and H3K27me3 ChIP-seq. (C) Heatmap showing expression change of all let-7 miRNAs in WT and *EZH2* -/- cells. Rows are ranked by low to high log2 fold change (WT/*EZH2* -/-).

**Supplemental Figure S11.**
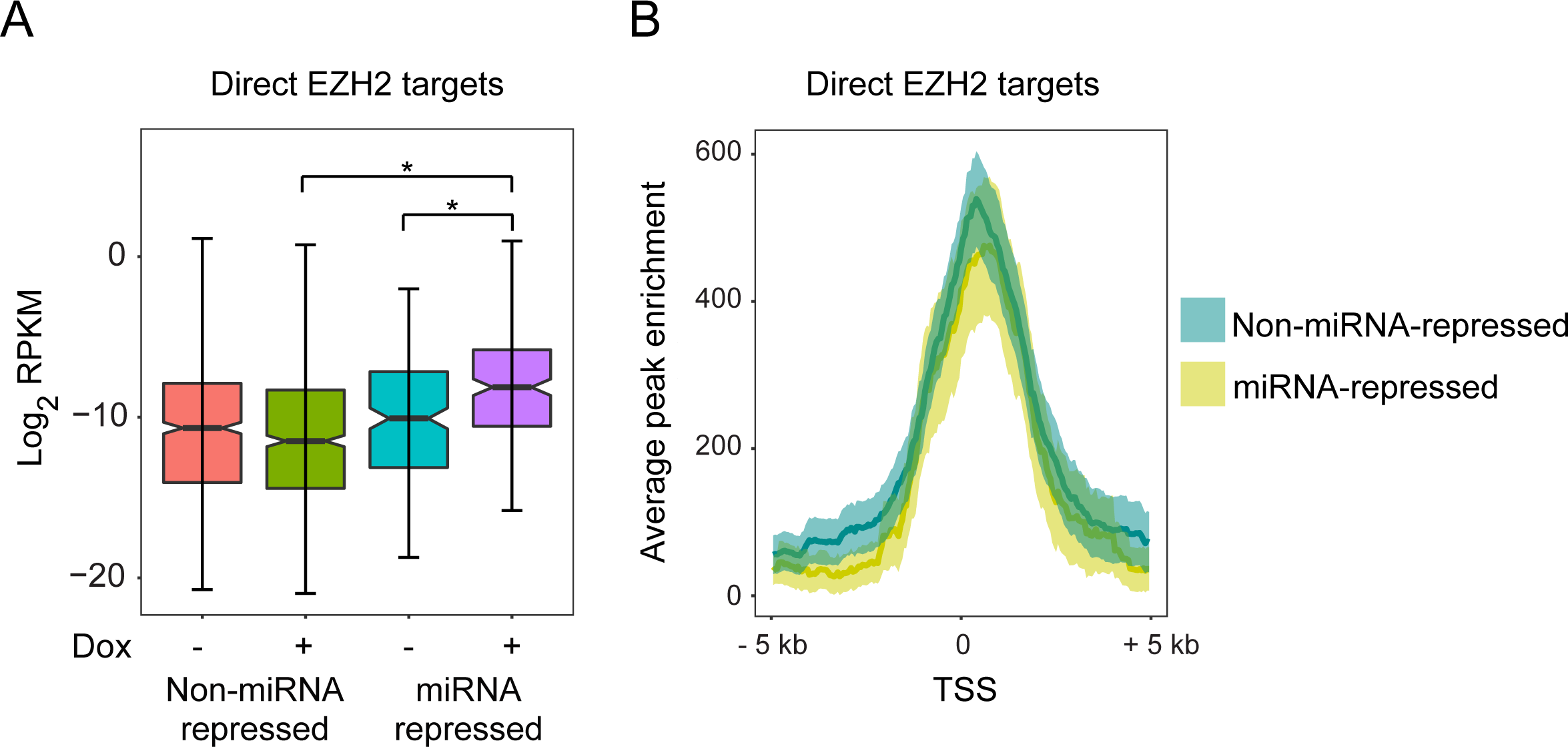
Genes co-repressed by *EZH2* and miRNAs are less tightly repressed by *EZH2* than genes silenced by *EZH2* alone. (A) Box plot showing average expression in Dox treated (miRNA depleted) and untreated WT cells for two sets of genes - PRC2-repressed genes that are not co-repressed by miRNAs (non-miRNA-repressed) and genes that are co-repressed by PRC2 and miRNAs (miRNA-repressed). (B) Normalized density of EZH2 ChIP enrichment scores in a 10 kb region spanning the TSS of PRC2 target genes that are either non-miRNA-repressed or miRNA-repressed.

## Abbreviations

PRC2: Polycomb repressive complex 2
TF: transcription factor
iCLIP: individual crosslinking and immunoprecipitation
UTR: untranslated region
RISC: RNA-induced silencing complex
Dox: doxycycline mESCs, mouse embryonic stem cells
GBM: Glioblastoma multiforme
TSS: Transcription start site(s).

